# Computational estimates of mechanical constraints on cell migration through the extracellular matrix

**DOI:** 10.1101/2020.01.19.912006

**Authors:** Ondrej Maxian, Alex Mogilner, Wanda Strychalski

**Affiliations:** Courant Institute of Mathematical Sciences, New York University, New York, NY, 10012 USA; Department of Biology, New York University, New York, NY 10012 USA; Department of Mathematics, Applied Mathematics and Statistics, Case Western Reserve University, Cleveland, OH 44106 USA

## Abstract

Cell migration through a three-dimensional (3D) extracellular matrix (ECM) underlies important physiological phenomena and is based on a variety of mechanical strategies depending on the cell type and the properties of the ECM. By using computer simulations, we investigate two such migration mechanisms – ‘push-pull’ (forming a finger-like protrusion, adhering to an ECM node, and pulling the cell body forward) and ‘rear-squeezing’ (pushing the cell body through the ECM by contracting the cell cortex and ECM at the cell rear). We present a computational model that accounts for both elastic deformation and forces of the ECM, an active cell cortex and nucleus, and for hydrodynamic forces and flow of the extracellular fluid, cytoplasm and nucleoplasm. We find that relations between three mechanical parameters – the cortex’s contractile force, nuclear elasticity and ECM rigidity – determine the effectiveness of cell migration through the dense ECM. The cell can migrate persistently even if its cortical contraction cannot deform a near-rigid ECM, but then the contraction of the cortex has to be able to sufficiently deform the nucleus. The cell can also migrate even if it fails to deform a stiff nucleus, but then it has to be able to sufficiently deform the ECM. Simulation results show that nuclear stiffness limits the cell migration more than the ECM rigidity. Simulations of the rear-squeezing mechanism of motility results in more robust migration with larger cell displacements than those with the push-pull mechanism over a range of parameter values.

**Author summary:** Computational simulations of models representing two different mechanisms of 3D cell migration in an extracellular matrix are presented. One mechanism represents a mesenchymal mode, characterized by finger-like actin protrusions, while the second mode is more amoeboid in that rear contraction of the cortex propels the cell forward. In both mechanisms, the cell generates a thin actin protrusion on the cortex that attaches to an ECM node. The cell is then either pulled (mesenchymal) or pushed (amoeboid) forward. Results show both mechanisms result in successful migration over a range of simulated parameter values as long as the contractile tension of the cortex exceeds either the nuclear stiffness or ECM stiffness, but not necessarily both. However, the distance traveled by the amoeboid migration mode is more robust to changes in parameter values, and is larger than in simulations of the mesenchymal mode. Additionally cells experience a favorable fluid pressure gradient when migrating in the amoeboid mode, and an adverse fluid pressure gradient in the mesenchymal mode.

## Introduction

The ability of cells to navigate complex three-dimensional (3D) extracellular matrix (ECM) is essential in the physiology of health and disease. One example of a process important for health is fibroblasts moving through the ECM to heal wounds [1]. On the other hand, one of the hallmarks of cancer is the migration of metastatic cancer cells across the ECM [2]. More often than not, cells move in cohesive groups, but many physiological phenomena involve single cell migration [3], which is our focus here. Understanding mechanics of this migration is a great present challenge, which can be met by combining cell biological, biophysical, and computational approaches [4].

There are a bewildering number of observed mechanical strategies of cell locomotion in 3D, reflecting the complexity and adaptability of the cell’s mechanical modules. The most frequently described migration modes are mesenchymal and amoeboid [5], but distinctions between these two modes are not clear-cut. In mesenchymal mode, cells are elongated and polarized, with protrusion activity at the short front, retraction activity at the short rear and opposite end, and tensed long sides. In this mode, integrin-dependent adhesions are distributed all over the cell surface and are crucial for migration, as inhibition of integrin stops the motion of the cells [6]. Cells protrude the front, form a pseudopod, attach it firmly to ECM fibers, and generate contraction within the pseudopod [7] and several microns behind its tip [8]. The pseudopod can be a cylindrical lobopod in a stiff ECM or a branched, finger-like lamellipodium in a soft matrix [6]. After the protrusion and contraction are generated, a continuous release of adhesions at the rear results in translocation of the cell body forward.

In amoeboid mode, cells are less polarized and have a more rounded shape. They have a more uniform distribution of cytoskeletal structures and/or membrane blebs around the periphery [5, 6] and migrate by squeezing through the pores of the ECM. One of the examples of the amoeboid migration is exhibited by epithelial cells in 3D collagen matrices [9], with observed nuclei leading the cell front, with contractile cell body trailing behind, and with actomyosin contraction propelling the nuclei forward, driving the migration of these cells. It appears from the images in [9] that the cells generate ECM deformations, which then propel the cells forward because the ECM is elastic and its deformations store elastic energy that can be released to propel the cell [10]. Another example of generating the ECM deformations and then harnessing them to create locomotion was recently reported in [11].

The modes of migration are malleable: cells are able to switch from one mode to another, depending on the physical and geometric properties of the cell and ECM [5, 6, 12]. What unites these modes is that cell migration in ECM depends crucially on myosin-powered contraction [4], which is not the case for cell migration on flat 2D surfaces [13]. The reason is that in 2D the cell’s largest organelle, the nucleus, is riding effortlessly atop the actomyosin locomotory network [14], while inside the ECM the problem of moving the bulky nucleus through the matrix becomes the center of the cell’s mechanical effort [4, 15]. Recent work indicates that the steric resistance the 3D matrix presents against the forward propulsion of the nucleus is a universal constraint for 3D cell migration [16–18], and myosin-generated forces are critical for overcoming this resistance.

Computational modelling is a valuable complement to experiments in understanding the complex cell migration mechanics. In 2005, Zaman et al. proposed one of the first, highly simplified, force balance models for 3D cell migration, which showed that the migration progress depends on adhesion strength and ECM mechanical and geometric properties [19]. One of the subsequent models included a continuum approach, in which each modelled cell was simplified as a self-protrusive 3D elastic unit interacting with an elastic ECM through detachable bonds [20]. Other, highly sophisticated, continuous mechanical models focused on the cell shape, rather than its interactions with deformable environment [21–23]. A few detailed, agent-based models of migrating cells immersed in a deformable ECM included models of the ECM, cell cortex, and membrane using networks of viscoelastic links [24, 25]. Another, very recent, effort investigated the influence of the flow of interstitial fluid on the cell migration through the ECM [10].

A few models recently started to investigate specifically the influence of the nucleus on 3D cell migration. Sakamoto et al. [26] proposed a computational model that took into account the viscoelastic property of the cell body. By using a finite-element method and prescribing cyclic protrusion of the leading edge of the cell, the authors predicted that the mesenchymal-to-amoeboid transition is caused by a reduced adhesion and an increased switching frequency between protrusion and contraction. A 2D mechanical model to simulate the migration of a HeLa cell with a large deformable cell body through a micro-channel was proposed in [22]. A very detailed model of a glioma cell represented by two elastic circular curves, an inner curve corresponding to the nucleus of the cell and an outer curve corresponding to the cell basal membrane, and explicit contractile myosin dynamics was proposed in [27]. In this model, migration of the glioma cell through a tissue made of normal cell represented by additional elastic curves was simulated with the immersed boundary method, with viscous fluid mediating interactions of the membranes and cortices. Last, but not least, a comprehensive mathematical model based on an energy minimization approach, was used to investigate cell movements inside a channel composed of ECM [28]. Simulations of this model reproduced deformations of the initially spherical nucleus treated as an elastic solid body into an elongated shape able to squeeze through the channel.

These models made inroads into the general question of what the mechanical constraints imposed by squeezing the nucleus through the ECM are on cell migration, yet the following specific questions remain open: Given nuclear and ECM mechanical properties, how strong should the myosin-powered contraction be to generate persistent cell locomotion? Could one or the other locomotory mode have a mechanical advantage? What are the migration mechanics if the nucleus is much stiffer than the ECM, or vice versa? What are the mechanical roles of the nucleoplasm, cytoplasm and interstitial fluid?

In this study, we model the cell as two elastic contours – one representing the nucleus, another, enveloping the nucleus, is the cell cortex. The cortex also has contractile and protruding activities. We model the ECM as an elastic network. Viscous fluids permeate the ECM (interstitial fluid) and fill the spaces inside the nucleus (nucleoplasm) and between the cortex and nucleus (cytoplasm). We model two mechanical strategies of cell migration: in one, resembling mesenchymal mode, the cortex makes a protrusion, the tip of which adheres to an ECM node and then global cortex contraction pulls the cell forward. In another, resembling one of the amoeboid modes, the cortex makes two attachments to ECM nodes and contracts only at one side, pushing the nucleus at that side through the ECM gaps at the opposite side. In order to simulate the coupled fluid-structure interaction problem, the model is formulated using the method of regularized Stokeslets [29]. In this method, the main force balance includes the elastic and contractile forces from the cell and ECM along with viscous fluid stresses. The fluid velocity computed by the method determines the dynamics of cell migration. We find that the migration is successful if the contractile tension of the cell cortex is greater than the characteristic force of either the nucleus or the ECM deformation, but not necessarily both. Simulations reproduce the observed amoeboid and mesenchymal morphodynamics and predict that the amoeboid mode overall performs better mechanically than the mesenchymal mode.

## Materials and methods

### Qualitative model description

To avoid the computational expense of 3D simulations, we consider a planar cross section of the cell and a cross section of the ECM in the same plane around the cell. One way to interpret the model’s geometry is to picture a cell extending a great distance in the direction perpendicular to the plane in which the structures and movements are explicitly simulated, and to assume that both the cell, and the ECM, and the fluid flow are homogeneous in that direction, so that all non-trivial effects occur in the 2D plane. Another approximation corresponds to considering an axially symmetric cell embedded into an axially symmetric ECM, and to neglecting geometric effects of the polar coordinate system. These approximations are not rigorous, so the model is, strictly speaking, 2D, but it captures the essential 3D effect of squeezing the deformable cell through the deformable ECM.

In short (mathematical and computational details are described in the following subsections and in the supporting text), the simulated cell consists of the elastic and protrusive-contractile contour, representing the cell’s actomyosin cortex and membrane. Inside the cell is a deformable nucleus represented by the elastic contour. Importantly, both the cell and nucleus are filled with fluid, with constant volumes (areas) of the nucleoplasm inside the nucleus and constant volume of the cytosol between the nucleus and cortex. The fluids are incompressible and viscous. Both cell and nuclear membranes are assumed impermeable to fluids for simplicity. The cell is embedded into an ECM represented by a 2D elastic node-spring network (shown in Fig. 1). Computationally, both the cell cortex and nuclear envelope, and ECM, are represented by 2D node-spring networks, similar to previous studies [25, 30].

**Fig 1.**
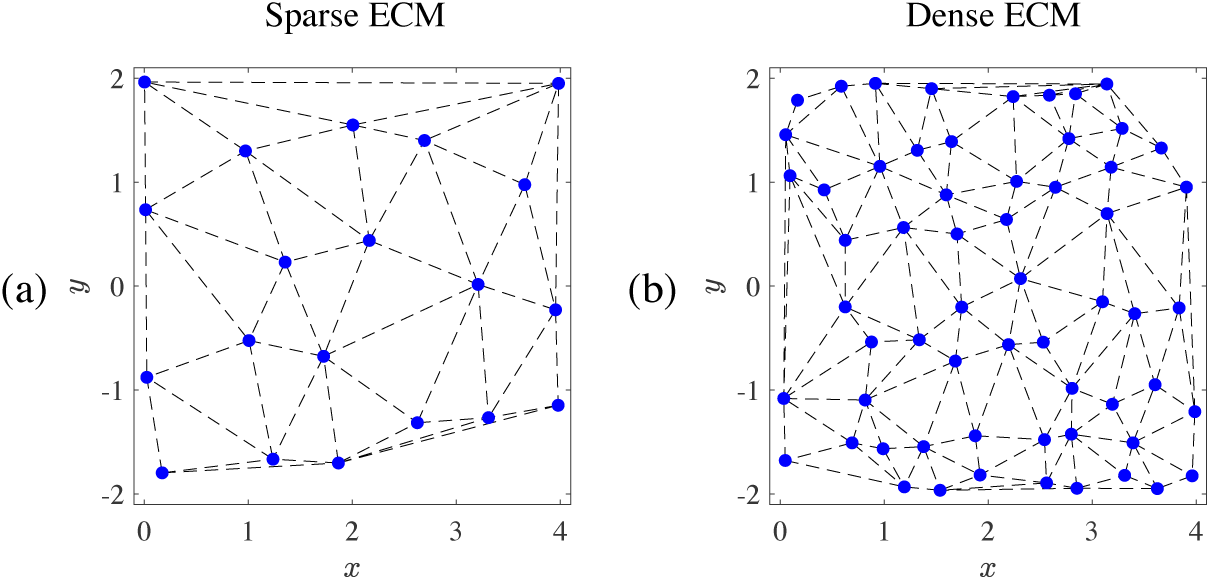
Sample ECMs used for cell motility simulations. The ECMs are generated on the box (*x, y*) ∈ [0, 4] × [−2, 2] (lengthscale units are in cell diameter). Blue points show the network, and black dotted lines show the spring connections The sparse network in (a) has 20 nodes, and the dense network in (b) has 60 nodes.

The ECM and cell cortex are immersed in the incompressible interstitial fluid. The viscosities of the nucleoplasm, cytoplasm and interstitial fluid are the same, for simplicity. The fundamental force balance in the model is implemented as follows: a net elastic/protrusive/contractile force at each node of the nucleus, cortex and ECM is applied to the fluid. The fluid’s velocity and pressure are then calculated by solving the Stokes’ equations using the method of regularized Stokeslets [29]. The method relies on the linearity of Stokes’ equations and involves using analytical formulas to compute the velocity and pressure distribution generated by a point-like force, then integrating over the material nodes where the forces are applied. Because all of the nodes move with the computed fluid velocity, the ECM nodes do not penetrate the cortex, and the nodes of the cortex do not penetrate the nucleus. Hence, there is no need to include repulsive forces in between different components.

We model two motile strategies. One of them loosely resembles the mesenchymal locomotion, during which cells migrate by forming a long, finger-like protrusion from the cell body into the ECM and generate traction forces behind the tip of the protrusion [8]. In the model, the cell first develops a narrowly focused outward-pushing force generating a finger-like protrusion that adheres to an ECM node. Then, global contraction of the cortex pulls the nucleus and cortex forward, ending the motile cycle. Fig. 2 I shows a schematic of this mode.

**Fig 2.**
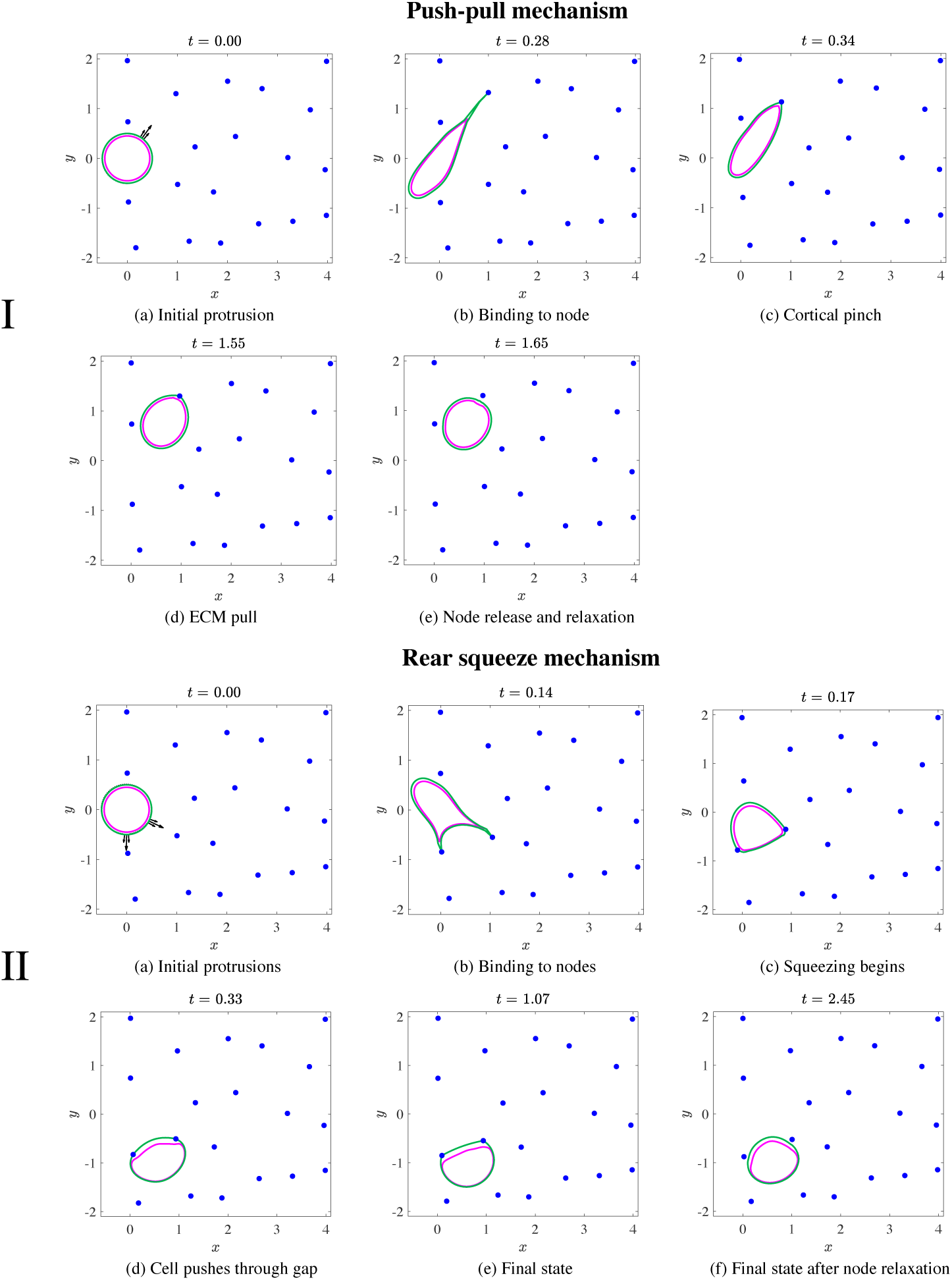
Stages of motility for the push-pull and rear-squeezing mechanisms of motility. The position of cortex (green), nucleus (magenta), and ECM nodes (blue), and applied forces (black arrows) are shown at several times values increasing from left to right and up to down. For panels in I, a random protrusion appears on the cortex (a) and extends until it comes into contact with an ECM node in (b). The cortex then becomes extremely stiff, which initially pulls the ECM node towards the cell in (c). Forces due to ECM elasticity eventually balance cellular forces to result in translocation of the entire cell (d). After coming to rest very close to the node, the cell releases the node and the node moves away. The system is then allowed to re-equilibrate, with the final state of one motility cycle shown in (e). Panels in II show the stages of motility when the cell uses rear contraction. Two protrusions form on the front (right) edge of the cell cortex in (a), then extend until both come into contact with ECM nodes in (b). The cortex is then separated into two regions. The leading edge region between the attachments remains loose, while the longer rear region behind the attachments contracts. Squeezing of the cell through the ECM gap occurs as the rear cortex contracts (c-e). The cell passes through the gap between the two nodes in (d), then comes to rest (e) when the region of the cortex between the two nodes is short and stiff. Next, the cell releases the ECM nodes, which move back toward their original positions in the final resting state (f), and the cortex relaxes. This completes one motility cycle.

Another motile strategy loosely resembles one of the amoeboid modes, in which the cortex first makes two protrusions that attach to two respective ECM nodes, followed by contraction of a half of the cortex behind the nucleus. This cortex contraction squeezes the nucleus forward through the ECM network [9, 16, 31]. Fig. 2 II shows a schematic of this mode.

We now describe the model equations governing mechanics of the ECM, cortex and nucleus, as well as the dynamic processes of the two motility mechanisms. The fluid mechanics that couple the solid deformations of the ECM, cortex and nucleus to the fluid are described in Section S1 of the supporting text.

### Extracellular matrix

The ECM in the model is composed of elastic fibers that are linked by virtual springs. We use the term “virtual” to indicate that the springs connecting the nodes do not sterically inhibit cell migration. Rather, we assume that cells interact only with the ECM nodes, and that the springs between the ECM nodes only provide restoring forces to the nodes. Thus, we treat the ECM simply as a collection of point-nodes. Suppose that there are *n* nodes in the ECM, then the ECM lattice is formed by triangulating these points. Any two nodes that share an edge (in the triangulation) are linked together by a virtual spring. Suppose that *N* (*i*) is the set of neighbors for the *i*-th fiber. Then the force on the ECM can be computed as

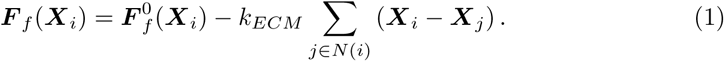

In Eq. (1), 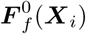 is the ‘pinning-down’ force that ensures the fibers are motionless at the beginning of the simulation (*t* = 0). Since we construct random lattices, there will be a net force initially on each fiber in the absence of 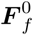 simply because the points are not located on a regularly spaced mesh (see Fig. 1). The addition of 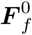 stabilizes the lattice and ensures that the fibers do not just collapse onto each other during a dynamic simulation (one can think of 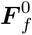 as mixture of an “anchor” for the whole lattice and effect of the springs’ resting lengths). The second term in Eq. (1) describes the sum of elastic contributions of the springs with stiffness *k*_*ECM*_ that connect fiber *i* to its neighbors.

We normalize lengths and distances so that the cell has diameter 1. The ECM nodes are therefore generated on the two dimensional box (*x, y*) ∈ [0, 4] × [−2, 2] (excluding a region near the origin where the cell is initially placed). Fig. 1 shows the ECMs used in our numerical experiments. We use 20 nodes for a sparse ECM and 60 nodes for a dense ECM. The black dotted lines show the elastic lattice that connects the nodes. Fig. 1 shows that the 20 and 60 node ECMs have average fiber spacings of about 1.5 and 0.5 respectively. This means that the ECM node spacing is either larger (for a sparse ECM) or smaller (for a dense ECM) than the cell diameter.

### Cell cortex and nucleus

The cell cortex is represented as an elastic contour, composed of discrete springs with a stiffness and rest length, that surrounds the nucleus, so that there is a small volume (area in 2D) of the cytoplasmic fluid between the nucleus and cortex. The net force in the cell cortex is a combination of passive elastic force, contractile force and protrusion force. We model the contractile force as an active contractile force that stems from stiffening the cortex springs, and, in one of the mechanisms, changing their rest length. Thus, mathematically, we can combine the passive elastic and contractile forces into the net elastic force, and to present the total force in the form:

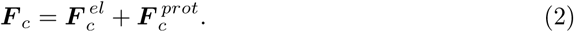

Elastic forces on the cortex are defined as follows:

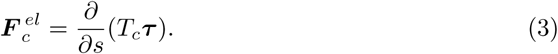

Here

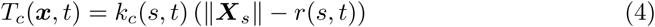

is the cortex tension, and ***X***_*s*_ is the non-dimensional tangent vector in reference arc length coordinates *s*. To account for contractile forces, the stiffness *k*_*c*_ of the contour springs can vary in time and with arc length parameter *s*. Similarly, the rest length *r*(*s, t*) can vary in space and time. In the relaxed state, *r* = 1. We note here that the central difference approximation to the *s* derivative ensures that the force 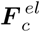 sums to zero, even when functions *k*_*c*_(*s, t*) and *r*(*s, t*) are not continuous functions of variable *s*.

The cortex contour is discretized by *N*_*c*_ points/nodes. The cortex generates a protrusion by first selecting a point on its perimeter, denoted by ***X***_0_. The protrusion force, centered on the point ***X***_0_, is in the locally normal direction to the cortex and has strength *f*_0_ at ***X***_0_ and strength *f*_0_*/*2 at the two points adjacent to ***X***_0_. The forces on these three points, directed outward, are balanced by the inward force spread over the other *N*_*c*_ − 3 points. The magnitude of the inward force is adjusted so that the total protrusion force is zero, and the cell does not move by applying a net force on the fluid. The value for *f*_0_ is chosen to be above a certain threshold to overcome the elastic resistance of the cell. Our choice of *f*_0_ results in a fast protrusion that does not contribute significantly to the dynamics of a single cell migration cycle, which is determined by viscous, elastic and contractile deformations. Fig. 2 I(a) shows the spatial profile of the force (in black) that generate the protrusion on the cortex (in green) in the 20 node ECM. Fig. 2 I(b) shows how the protrusion expands.

The cell nucleus is modeled as a simple elastic contour. It is discretized with *N*_*n*_ points, and the elastic force is computed with Eq. (3) with *T* = *k*_*n*_(‖***X***_*s*_‖ − 1). For the nucleus, the stiffness *k*_*n*_ and rest length are constant throughout the simulation.

### Motility mechanisms

#### Mechanism 1: push-pull mesenchymal mode

In the first case, the cell generates one protrusion at a time on its front half (facing the right). We do not investigate the process of choosing the direction of protrusion, as we focus on the mechanics of the motility cycle. Thus, we choose a point ***X***_0_ on the right half of the cortex at random (i.e. from a uniform distribution) for the protrusion to form. The overall process is shown in Fig. 2 I (top two rows). The cortex, which is initially relaxed (small stiffness 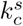) generates a protrusion (Fig. 2 I(a)) which extends until it comes into contact with an ECM node. Numerically, a contact occurs when the point at the tip of the protrusion comes within a small threshold from an ECM node. The tip of the protrusion then reaches out and attaches to the ECM node, as shown in Fig. 2 I(b).

Once the cortex attaches to the ECM node, it immediately stiffens globally. Stiffening of the cortex is modeled by increasing the parameter for cortical tension *k*_*c*_ to a value 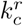, which is varied to investigate different motile regimes. The cortex then rapidly rounds to assume its circular resting configuration. In the case of a rigid ECM, the process results in the round cortex being pulled toward the attached ECM node. For the case of a softer ECM, the dynamics are more subtle. First, as shown in Fig. 2 I(c), the cortex assumes a quasi-circular shape by pulling the ECM node toward the cell. This inward pull continues until the elastic deformation force from the ECM balances the pulling force from cortical elasticity. The balancing of these opposing forces evolves, so that as the cortex becomes more circular, the ECM becomes less deformed, and the pulling force decreases. Eventually, the ECM node comes to rest near its initial position, as in Fig. 2 I(d). Once the velocities of the ECM and cortex nodes relax below a small threshold value, the cortex detaches from the ECM node. In order to physically uncouple the motion of the cortex and ECM node in the presence of the fluid, we move the ECM node a small threshold distance away from the cortex and wait for the system to equilibrate again (i.e. for the maximum velocity of the ECM and cortex nodes to relax below the small threshold value). The end result of this process is shown in Fig. 2 I(e). The cortex is then free to form another protrusion. We will refer to this entire process as the motility cycle.

We note that not all protrusions result in cellular motion. In fact, for sparse ECMs, some protrusions will not contact any nodes. For this reason, in mechanism 1 we automatically retract (by stiffening the cortex globally) any protrusions that exceed a length of four cortical radii, assuming that they were not able to find any nodes. We also refer to this case as a cycle, so that cycles can be either successful or unsuccessful.

#### Mechanism 2: Rear-squeezing mode

In this mechanism of motility, two protrusions are generated on the loose cell cortex. As in the push-pull mechanism, we choose the first protrusion location randomly on the front half of the cell. The second protrusion is generated at a location 30-45° apart from the first protrusion so that the protrusions are positioned to grow towards two adjacent ECM nodes. The resulting force distribution for these two protrusions is the superposition of two mechanism 1-type protrusions and is shown in Fig. 2 II(a). The protrusions expand until each contacts a node as in Fig. 2 II(b). Once contact occurs on both protrusions, the attachments between the protrusions and respective ECM nodes form, and the longer length of the cortex behind the two attachments becomes very stiff, with the stiffness increasing to some value 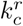 (varied in numerical experiments), while the non-dimensional discrete spring rest length decreases from *r* = 1 to *r* = 0.1. We have used the same parameter 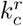 across both mechanisms to represent increased cortical tension, although this tension is only increased in part of the cell in mechanism 2. Also, note that decreasing the rest length to *r* = 0.1 is equivalent to adding an active tension (contraction) at the rear of the cell, and is required to generate enough tension to squeeze the cell through the ECM.

The shorter length, i.e. the leading edge at the right side, of the cortex between the two nodes remains relaxed (stiffness remains equal to 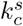, and *r* = 1). The contracting rear of the cortex squeezes the nucleus through the gap between the two attached ECM nodes by deforming both nucleus and ECM, resulting in cell propulsion. This process is shown in Figs. 2 II(c)-(e). Once the system comes to a resting state (all node velocities decrease below the threshold), both nodes are moved a small threshold distance away from the cell, and the system (cell and ECM nodes) is allowed to equilibrate until it comes to rest. The cell is then ready to form another protrusion (the final state is shown in Fig. 2 II(f)). This entire process defines one cycle of motility.

Two ECM contacts are required to move the cell, and protrusions sometimes fail to reach a node in this mechanism. As before, we retract protrusions that reach a certain length before hitting a node, and restart the process. In order to maintain the same total protrusive length across mechanisms, we stipulate the maximum length of each protrusion as 2 cortical radii, so that the maximum total protrusive length of 4 cortical radii is the same as in mechanism 1. Alternatively, if only one of the protrusions contacts a node and the other reaches its maximum length without contact, the entire system is contracted (by increasing globally the cell stiffness) until the system returns to rest and can generate new protrusions. Although the cell does not move in this case, this too defines a cycle, so that once again cycles can be successful or unsuccessful.

### Parameter values and numerical procedure

#### Physical parameters

Geometric parameters include the ECM mesh size and sizes of the cell cortex and nucleus, which are all of the same order of magnitude [5, 32]: cell diameter is of the order of 10 *µ*m, while the nuclear diameter is slightly smaller [33]. Openings in 3D extracellular environments range from 2 to 30 *µ*m in diameter [34]. Thus, we choose the cell cortex and nucleus diameters at rest to be equal to 10 and 9 *µ*m, respectively. The length is normalized in all simulations to make the cell diameter, 10 *µ*m, to be the unit of length. We vary the ECM density so that the average distance between the nearest ECM nodes is 1.5 or 0.5 length units, in the cases of low- and high-density ECM, respectively, so that the cell can move through the ECM undeformed in the low-density case, and has to deform significantly to squeeze through the ECM in the high-density case.

The issue of physical dimensions in a 2D model that includes interactions of solid and fluid structures is subtle. From the hydrodynamic part of the model (see Section S1 in the supporting text), it is clear that in the 2D method of regularized Stokeslets the force applied to the fluid at a point has the dimension of pN/*µ*m (see Eq. (S1.1): fluid velocity is in *µ*m/sec, viscosity is in Pa·s, hence the the force is in pN/*µ*m.) The interpretation is as follows: the 2D approximation is the planar cross-section of the 3D space, and the ‘per micron’ factor appears in the force because this force is ‘per unit length in the direction perpendicular to the 2D plane’. Simply speaking, to estimate the physical force in 3D space, one has to multiply the pN/*µ*m force in the 2D model by the characteristic cell size, 10 *µ*m. These considerations affect the choice of the mechanical model parameters.

There is, as expected, a significant variability in the reported values of the ECM, nucleus and cell cortex mechanical characteristics. In our 2D model, the ECM spring constant, *k*_*ECM*_, is measured in the units of pN/*µ*m^2^, so that when multiplied by the spring extension or shortening (in *µ*m), the force on the point-like node has unit of pN/*µ*m. The Young modulus of the collagen mesh and ECM can vary from 1 to hundreds of Pa [12, 35–37]. We choose ECM spring stiffness of *k*_*ECM*_ = 50 pN/*µ*m^2^, which corresponds to the Young modulus within the range of values reported in the literature. We keep this parameter the same in all simulations.

The elastic and contractile force densities per unit length of the respective contours in the cell cortex and nuclear envelope have dimensions pN/*µ*m^2^ in our 2D model, which, when multiplied by the characteristic distances between the discretization nodes in the contours, turn into forces with dimension pN/*µ*m applied to the fluid. These force densities are spatial derivatives of the tensions along the contours, so these tensions have dimensions of pN/*µ*m. Note that in Eq. (3), the arc length *s* is dimensional, in *µ*m, while in Eq. (4) the expression for strain in brackets is non-dimensional. The same considerations apply to the nuclear mechanics in the model. Thus, *k*_*c*_ and *k*_*n*_ are the dimensional proportionality coefficients in the cortex and nuclear contours’ tensions, respectively, in pN/*µ*m (the tensions itself are these proportionality coefficients times the non-dimensional strain). These proportionality coefficients are effectively spring constants for the respective contours, which correspond to respective Young moduli (measured in pN/*µ*m^2^ in 3D) after being divided by the characteristic cell size, 10 *µ*m.

The mechanical modulus of the nucleus can vary from tens of Pa [38] to hundreds of Pa [35, 39] to thousands of Pa [40], corresponding to values of *k*_*n*_ from units to hundreds of pN/*µ*m. In the simulations, we vary the nuclear stiffness in the range even wider than that reported, from 1 to 10000 pN/*µ*m, with 1 – 10 pN/*µ*m corresponding to ‘soft’, ∼ 100 pN/*µ*m corresponding to baseline, and 1000 – 10000 pN/*µ*m corresponding to ‘stiff’ nucleus. In the tables below, we report the nuclear stiffness 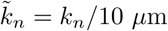, to make consistent comparison with the ECM stiffness.

The tension of the cell cortex (in units of pN/*µ*m, which directly corresponds to our parameters 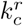 for a contracting cortex) was reported in the range from ∼ 100 pN/*µ*m [33] to ~ 1000 pN/*µ*m [41]. We use the value 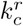 varying up to 1000 pN/*µ*m, except in one of the numerical experiments, where we simulate an exceedingly weak cortex with 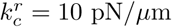 (in which case, the cortex is still contracting due to a decrease in its discrete spring rest length). For consistent comparison with the ECM stiffness, in the tables below we report 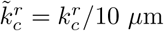. We use the value 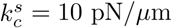 for the loose, relaxed cortex, which corresponds to 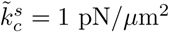. Further discussion of the mechanics of nucleus and cortex can be found in [17, 42].

The last physical parameter we have in the model is the fluid viscosity. The actual physical value does not matter for the simulations, because the viscosity simply determines the time scale. In the simulations, the viscosity is normalised to 1. Nevertheless, it is instructive to discuss the physical values of the viscosity of the nucleoplasm, cytoplasm and interstitial fluid that could be one to three, and even four orders of magnitude greater than of water [36, 37, 43–45]. In physical units, this means viscosity from 0.01 to 10 Pa·s. In the processes of the 3D cell migration, the timescale of viscous relaxation was observed to be in the order of tens of seconds [39]. In our simulations, characteristic net tensions of ∼ 10 pN/*µ*m (greater elastic and contractile forces largely equilibrate, so the net force is relatively small), characteristic sizes of 10 *µ*m and characteristic viscosity of 10 Pa·s result in ∼ 10 s time scale which compares well with observations reported in [39]. Of course, the additional processes of developing and relaxing protrusions, adhesion and contractions can add a significant time to this estimate; no wonder that 3D motile cycles were reported in the range of tens of minutes [46].

#### Numerical parameters

The numbers of grid points for both the cortex, *N*_*c*_, and nucleus, *N*_*n*_, contours in the simulations were varied depending on the motility mechanism and parameter set. In general, we set *N*_*c*_ = *N*_*n*_ = 50 − 80 for mechanism 1 and *N*_*c*_ = 80 − 300, *N*_*n*_ = 60 − 80 for mechanism 2. The number of cortex points is larger for mechanism 2 because the cell gets closer to the ECM nodes, which can leak inside the cell if the cortical boundary is insufficiently resolved. Even with *N*_*c*_ = 300 points there can still be minor blurring of the ECM/nuclear/cortical boundaries on the length scale of 0.1 *µ*m. This does not affect our conclusions here, as a test for a single cycle with *N*_*c*_ = 600 and *N*_*n*_ = 120 had a displacement that differed from the same cycle with *N*_*c*_ = 300 and *N*_*n*_ = 60, respectively, by < 2% (for both the smallest and largest nuclear stiffness used). The maximum stable time step is dictated by ECM stiffness and is 0.001 (in dimensionless time units) for *k*_*ECM*_ = 50 pN/*µ*m^2^. We decrease this timestep down to 0.0002 for a short period of time (𝒪(0.1) time units) when the cortex stiffens as it binds to the ECM nodes.

We note that mechanism 2 is much more challenging numerically than mechanism 1 because of the changing spring rest length *r*. Once the cortex binds to the ECM nodes, grid points behind the ECM nodes begin to pack very close together due to the shorter rest length. Meanwhile, points that span the front section of the cortex between the two ECM nodes grow apart quickly as the entire nucleus squeezes through the two nodes, with only a few grid points in front of it. The region in front of the two nodes therefore becomes under-resolved, and dynamic re-meshing is required to keep a stable configuration of the cortex and nucleus. This re-meshing takes place every 𝒪(100) time steps and only when the cell is attached to two ECM nodes. Because the nucleus interacts with the cortex via hydrodynamic forces, it too becomes under-resolved at the front, so the re-meshing is applied to both the cortex and the nucleus.

Our goal was to dynamically update the discretization so that the spacing between the grid points stays roughly constant through time. We accomplish this as follows. Given a current set of the grid points ***X***(*j*) with reference points ***X***^0^(*j*), we use the positions ***X***(*j*) to construct a cumulative arclength function, defined as *s*(*j*), where *j* is the point index. We then sample *s*(*j*) at equal lengths. Each of these lengths corresponds to a value of *j*. We denote the set of equally spaced arc lengths as *S*(*j*). Then the new point positions are given by ***X***(*S*(*j*)) and the corresponding new reference points are given by ***X***^0^(*S*(*j*)). We then update the positions of the grid points along with their reference configurations according to the new positions ***X***(*S*(*j*)) and ***X***^0^(*S*(*j*)), which can be found by linear interpolation (in general the values of *S*(*j*) are not taken at integer values of *j*). See Section S2 of the supporting text for more details about the re-meshing algorithm.

## Results

### Cells can use both mechanisms to move through a sparse ECM

Simulations of a cell moving through a sparse ECM (20 nodes; the mesh size is greater than the cell size) over one motility cycle are shown in Fig. 2. Additional time values are provided in Fig. S2 in the supporting text for a cell migrating with the push-pull mechanism and in supporting video S1. A cell migrating with a rear-squeezing mechanism in a sparse ECM is shown in supporting video S3. The figures and movies show that the cell is able to move through the ECM with minor deformation. In this regime, the limiting factor is not mechanics, but rather timing of developing protrusions and attachments and detachments to the ECM. Simulation data show that the cell can move without deforming its nucleus or the ECM. Therefore, it can move effectively regardless of the nuclear stiffness and contractile force. Therefore, we do not investigate the sparse ECM further and turn to the case of a dense ECM of 60 nodes (see Fig. 1(b)). Note that it was observed that in sparse matrices, cells use mechanisms entirely different from the ones we are considering, like wrapping tightly around one ECM fiber and moving along it [4, 47], which intuitively is a much more logical way to move than trying to search for far away ECM nodes.

### Mechanism 1 performs best for high tension and soft nucleus

We next consider a denser ECM (60 nodes; the ECM mesh size is roughly half the cell size) and a few characteristic regimes of the parameters and analyze the cellular motion therein. There are three characteristic force scales in the process: (1) *T*, the characteristic contractile tension in the cortex, the order of magnitude of which is given by the parameter 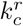. (2) *E*, the characteristic tension of the ECM deformed by the nucleus squeezing through it. This tension is of the order of the ECM spring coefficient *k*_*ECM*_ multiplied by the cell size minus the average ECM mesh size. (3) *N*, the characteristic tension in the deformed nuclear envelope, which is of the order of the the nuclear stiffness *k*_*n*_. In this section we consider all qualitatively different relations between these three forces. Table 1 gives the parameter choices for each of these regimes.

**Table 1.** Parameter values for the different forcing regimes for both migration mechanisms.

| Push-pull mechanism |  |  | Rear squeeze mechanism |  |  |
| --- | --- | --- | --- | --- | --- |
| Regime | $\tilde{k}_n$ (pN/$\mu\text{m}^2$) | $\tilde{k}_c^r$ (pN/$\mu\text{m}^2$) | Regime | $\tilde{k}_n$ (pN/$\mu\text{m}^2$) | $\tilde{k}_c^r$ (pN/$\mu\text{m}^2$) |
| $E, N > T$ | 100 | 1 | $E > T, N > T$ | 10 | 1 |
| $T > E > N$ | 1 | 100 | $T > E > N$ | 0.1 | 100 |
| $T > N > E$ | 10 | 100 | $T > N > E$ | 10 | 100 |
| $E > T > N$ | 1 | 1 | $E > T > N$ | 0.1 | 10 |
| $N > T > E$ | 100 | 50 | $N > T > E$ | 1000 | 100 |

Additional simulation information: there are 60 ECM nodes and $k_{ECM} = 50$ pN/ $\mu\text{m}^2$ .

We begin by discussing a regime in which the cell cannot move. This regime is actually composed of two orderings of the forces: *E* > *N* > *T* and *N* > *E* > *T*. In either case, both the force required to deform the nucleus and the force required to deform the ECM are greater than the contractile force. In this case, the position of the cortex, nucleus, and ECM at two time values are shown in Fig. 3 I(a). Additional simulation time values are shown in Fig. S3 of the supporting text and in supporting video S2. Results show that the cell becomes “stuck” in the ECM and is unable to pass through the small gaps in the dense ECM.

**Fig 3.**
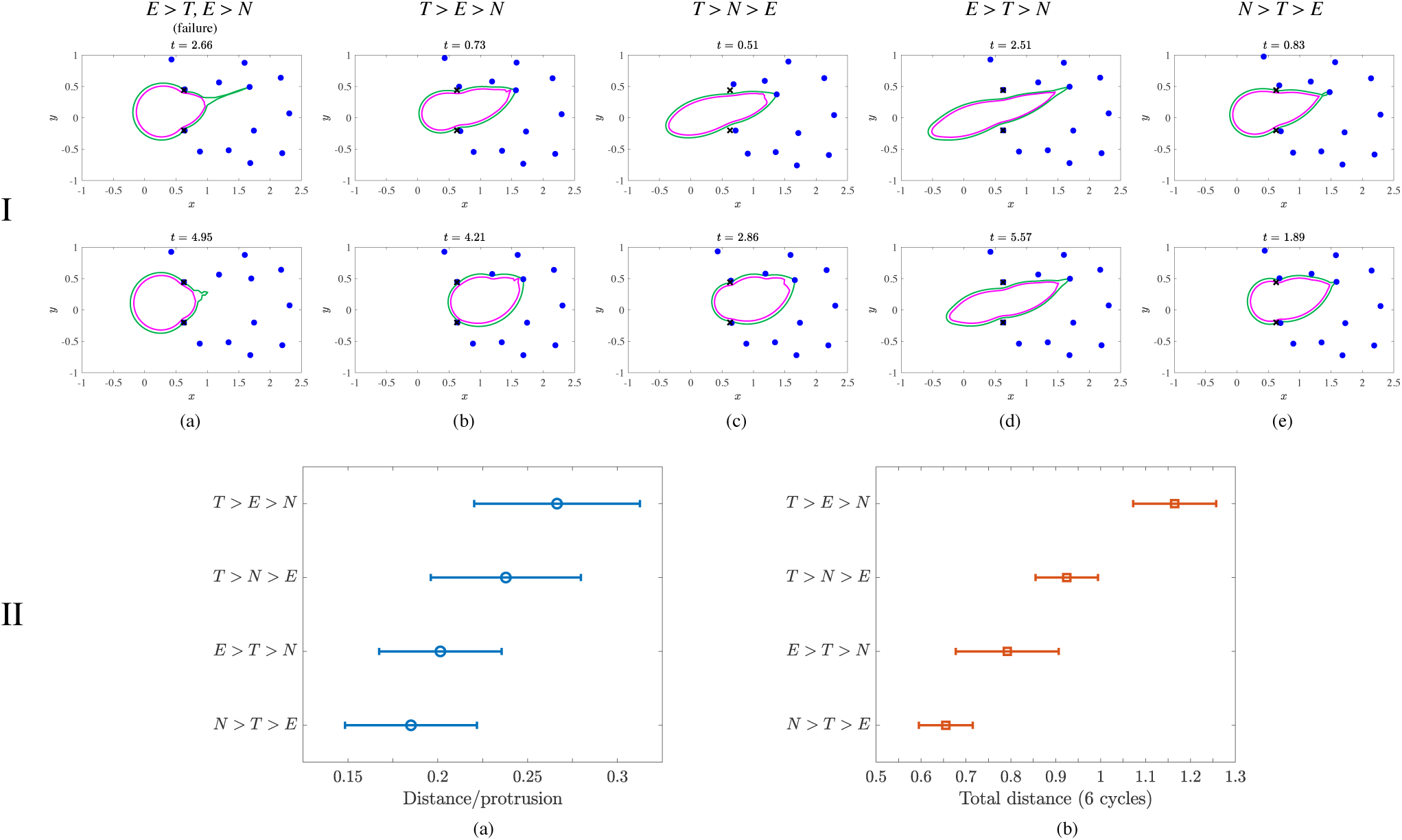
Results from simulating cell migration using the push-pull mechanism. Row I shows the position of the cortex (green), nucleus (magenta), ECM nodes (blue), and initial position of the ECM (black ×‘s) over a range of parameter values listed in Table 1. Simulation time values are located above each panel. Row II shows normalized distance of the cell from its initial location (distance is normalized by the cell diameter at rest). The average distance traveled over one cycle is shown in (a). The average distance per simulation (total displacement over 6 cycles) for each of the parameter regimes simulated is shown in (b). The data show a stronger contractile force *T* combined with a weaker nuclear force *N* leads to maximum displacement for both measurements. Error bars are a single standard error in the mean in each direction.

We next consider the case *T* > *E* > *N*, when the contractile force is the largest and the nuclear force is the weakest. The position of the cell and ECM from a simulation is graphed in Fig. 3 I(b) with more time values provided in the supporting text, Fig. S4. Data show that the nucleus is highly deformable and can easily slide through the gaps of the ECM nodes for these parameter values. The strong contractile force contracts the cell rear, so that the entire cell is able to slide through the ECM efficiently.

When the force required to deform the nucleus increases, the cell moves by deforming the ECM. Cell and ECM position from simulations in the parameter regime *T* > *N* > *E* are shown in Figs. 3 I(c) and S5 (supporting text). The difference between the regimes *T* > *N* > *E* and *T* > *E* > *N* can be seen in Fig. 3 (compare (c) to (b)), where the ECM nodes are further displaced from their original locations (shown as black ×’s) in the former regime than the latter.

In the parameter regime *E* > *T* > *N*, the ECM stiffness is the largest force relative to the other parameters, and the cell must move by deforming its nucleus. Simulation data in Fig. 3 I(d) shows that the ECM nodes do not move from their original positions and the ECM is essentially rigid. Additional time values for the position of the cortex, nucleus, and ECM nodes are provided in Fig. S6 (supporting text). Position data show that the nucleus is able to slide through the gaps, but the relatively weaker contractile force is sometimes unable to efficiently contract the rear of the cell. Several motility cycles are necessary for the entire cell to squeeze through a gap in the ECM. Fig. 3 shows that this regime gives the longest and thinnest cell shape, as it must pass through gaps without deforming the stiff ECM. However, simulation data in Fig. S6 show that the cell can easily deform to slide through the ECM nodes.

For the parameter regime *N* > *T* > *E*, the nuclear force is larger than both *T* and *E* so that the cell migrates by deforming the ECM, as shown in Fig. S7 in the supporting text. In fact, the maximum ECM displacement for data in Fig. S7 is 50% larger than for the parameter regime *T* > *N* > *E* (data shown in Fig. S5). Fig. 3 shows that this regime has the roundest cell shape as it migrates through the matrix when compared to other successful parameter regimes. The relatively large value for stiffness of the nucleus makes it difficult for the cell to move through a narrow gap when there is not enough force to move ECM nodes. The rear of the cell can become lodged in the ECM, as shown in Fig. 3 I(e), and overall this parameter regime is not very effective for migration.

#### Quantitative comparison of parameter regimes

We performed 8 simulations for the parameter regimes listed in Table 1, where the cell undergoes 6 protrusion cycles (protrude, bind, release, relax, repeat) per simulation (the total number of protrusions is 48). We measure the net distance moved per cycle and the net distance moved at the end of the simulation (total displacement) for the different parameter regimes. Distances are measured by tracking the center of mass of the nucleus (mean coordinates of the discrete points around the nuclear contour).

Fig. 3 II shows that the regime *T* > *E* > *N* provides the optimal motion for both an average cycle (Fig. 3 II(a)) and entire simulation (Fig. 3 II(b)). In both cases, *T* > *N* > *E* provides the second-best motility, where the tension is kept the same but the nucleus is stiffer than the ECM. We can interpret the decrease in distance traveled from *T* > *E* > *N* to *T* > *N* > *E* as follows. When the ECM is softer, it deforms more easily. At the end of a motility cycle, the ECM nodes begin to return to their initial position, which can inhibit the cell’s ability to migrate through the gap (see the two frames shown in Fig. 3 I(c)). The elastic restoring force from the ECM induces a small flow that results in the cell moving slightly backwards, negating some of its forward progression. On the other hand, when the nucleus is soft, the cortical contraction results in nuclear deformation, and the cell is able to squeeze through the ECM gap. Less effective parameter regimes are characterized by relatively reduced cortical contraction. When the cell cannot deform the stiff ECM, *E* > *T* > *N*, the relatively soft nucleus can still be deformed enough so that the cell can be pulled through the gap. The least effective regime is when *N* > *T* > *E*. In this case, the nucleus is the stiffest and the contractile force on the cortex is not large enough to deform the nucleus. The cortical contraction still can deform the ECM, but forces due to the elastic restoring force of the ECM inhibit the cell’s progress.

### Mechanism 2 results in robust cell migration

We analyze five different parameter regimes representing combinations of three characteristic forces for cells migrating using mechanism 2. Parameter values and combinations are listed in Table 1. Simulation results show that for a sparse ECM, a cell is able to successfully migrate as long as it can find two nodes (see supporting video S3; this point is further elaborated on in the discussion section). The position of the membrane, cortex, and ECM during a simulation of one motility cycle is shown in Fig. 2 II.

The cell is unable to migrate in the parameter regime *E* > *T* and *N* > *T*. The position of the cortex, nucleus, and ECM at two time values from a simulation is shown in Fig. 4 I(a) with several more time values provided in Fig. S8. A movie showing the position of the cell and ECM during a simulation is provided in the supporting video S4. The data show that when the cortex contraction is too weak to deform either nucleus or ECM, the cell is not able to squeeze through the ECM gap.

**Fig 4.**
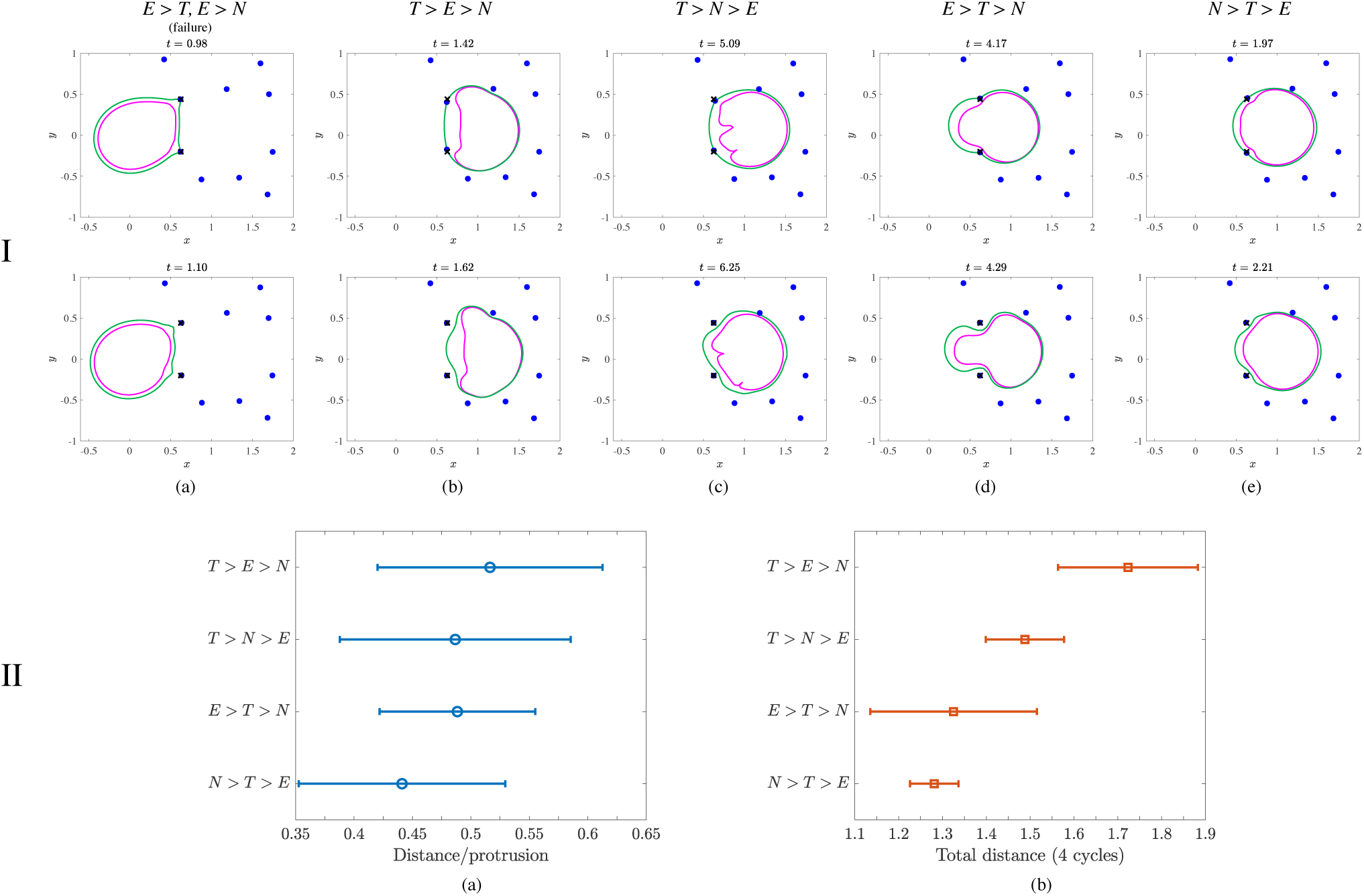
Results from simulating cell migration using the rear squeezing mechanism. Row I shows the position of the cortex (green), nucleus (magenta), ECM nodes (blue), and initial position of the ECM (black ×‘s) over a range of parameter values listed in Table 1. Simulation times values are located above each panel. Row II shows normalized distance traveled by the cell from its initial location (distance is normalized by the cell diameter at rest). The average distance moved over one cycle is shown in (a). The average distance per simulation (total displacement over 4 cycles) for each of the parameter regimes simulated is shown in (b). The data show that a relatively weak nuclear force *N* leads to maximum displacement for both measurements. Error bars are a single standard error in the mean in each direction.

When the contractile tension on the cortex is increased, for example in the parameter regime *T* > *E* > *N*, the cell is easily able to pass through the ECM network. Cell and ECM position during migration is graphed in Fig. 4 I(b) with more time values provided in Fig. S9. The cortex contracts at the rear and squeezes the nucleus through the gap between ECM nodes. However, when cortical tension remains high, but the nucleus is stiff relative to the ECM (for the parameter regime *T* > *N* > *E*), the cell does not completely pass through the ECM after one motility cycle (compare Fig. 4 I(b) to I(c)). The nucleus in Fig. 4 I(c) (top panel) appears to “buckle” under high cortical tension. More time values from a simulation in the parameter regime *T* > *N* > *E* are shown in Fig. S10. After the cortex detaches from the nodes, the nucleus rounds because of decreased tension at the cell rear. As the nucleus rounds, the cell slightly retracts due to the induced flow caused by the dynamic elastic forces on the nucleus. The entire process is illustrated in the supporting video S5. Subsequent cycles result in successful migration of the cell completely through the ECM gap, but the overall amount of motion apppears reduced compared to simulations in the parameter regime *T* > *E* > *N*.

In the parameter regime *E* > *T* > *N*, the nucleus is more deformable. Fig. 4 I(d) shows a highly deformed nucleus as the cell migrates through a gap (additional time values graphed in Fig. S11, supporting text). However, the cortex and nucleus do not completely clear the gap after one motilty cycle because contraction on the cortex is not larger enough relative to ECM stiffness.

Finally, in the parameter regime *N* > *T* > *E*, the nuclear stiffness is large relative to other parameters, and the cell has enough tension to deform the ECM. Fig. 4 I(e) shows an overall round cell during a successful motility cycle (additional simulation time values provided in S12 in the supporting text). The cell migrates by pushing the relatively soft ECM nodes out of its way. However, the cell’s passage through the gap is incomplete, for the same reason as explained in the parameter regime *T* > *N* > *E*; once the cell detaches, the nucleus rounds and a small opposing flow is induced that impedes the cell’s progress. Subsequent motility cycles result in the complete passage of the cell through the ECM gap. Note the increased stiffness of the nucleus prevents the buckling seen in Fig. 4 I(c).

#### Quantitative comparison of parameter regimes

As for mechanism 1, we perform 8 simulations, where each simulation the cell goes through 4 cycles of mechanism 2 (we perform 4 cycles because the distance per cycle is larger than in mechanism 1). We again measure the net distance migrated per cycle and the net distance moved (total displacement) at the end of the simulation across the different parameter regimes listed in Table 1. There are a total of 32 total cycles analyzed. Results are graphed in Fig. 4 II along with the standard error on the mean.

Simulations with parameter regimes where the nuclear stiffness is relatively small result in the largest distance traveled. Similar to results from the first mechanism, when the ECM is the softest element, the nodes partially recoil at the end of a motility cycle. The flow induced by the elastic deformation of the ECM causes the cell to partially retract. When the nucleus is the softest element, the entire cell can be pushed through the stiff matrix. Overall, we observe that increased cortical contraction results in larger distances traveled during migration for this mechanism.

Comparing quantitative results from simulations where the cell uses mechanism 1 (push-pull) to mechanism 2 (rear-squeezing), mechanism 2 results in relatively increased distance traveled over a range of parameter values (compare relative distance values in Fig. 3 II to Fig. 4 II). For example, in mechanism 1, the least effective parameter regime gives a displacement that is ≈ 70% of the maximum (most effective regime) for a single protrusion and ≈ 55% of the maximum for a simulation of 6 protrusions. Thus the spread in the data is almost 50%. Meanwhile, in mechanism 2, these quantities are ≈ 85% for a single protrusion and ≈ 75% for a cycle of 4 protrusions. Thus the spread in the data for mechanism 2 is approximately half of what it is in mechanism 1. For this reason, we describe mechanism 2 as more robust than mechanism 1. Furthermore, in all regimes, the displacements of the cell are greater for mechanism 2 than for mechanism 1 (discussed in the following section). That said, we note that a cell migrating using mechanism 2 experiences increased cortical tension compared to mechanism 1 because the rest length of the cortex at the back of the cell in this mechanism is decreased by 90%.

### Mechanism 2 is aided by hydrodynamics

Pressure during a motility cycle for both mechanisms of motility during comparable stages is shown in Fig. 5 when cortical tension is the dominant force (for the parameter regime *T* > *E* > *N* listed in Table 1). The cell in Fig. 5(a) is in the ‘ECM pull’ stage (see Fig. 2 I(d)), and the cell in Fig. 5(b) is in the ‘squeezing begins’ stage in Fig. 2 II(c). When there is a favorable pressure gradient, fluid flows from regions of high to low pressure, but fluid flow can still occur from low to high pressure. One example occurs in flow past an airfoil [48]. Fig. 5(b) shows there is a favorable pressure gradient with high pressure in the cell rear along with low pressure in the front of the cell for mechanism 2. The pressure gradient, which results from compression of the fluid at the rear of the cortex in mechanism 2, induces an additional fluid velocity in the direction of migration.

**Fig 5.**
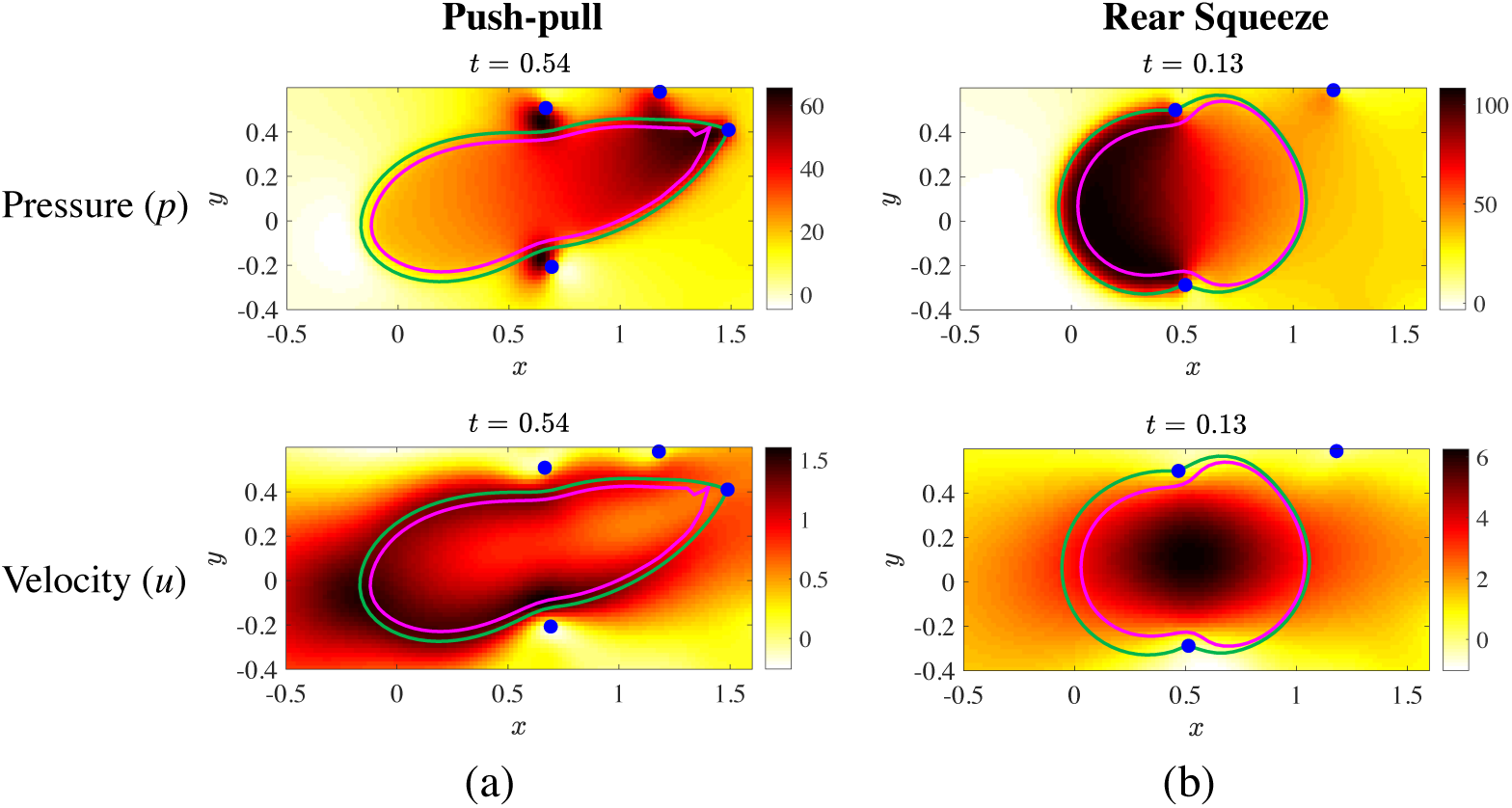
Pressure and horizontal component of velocity for a for a cell using (a) the push-pull and (b) rear-squeeze mechanisms of motility. The position of the cortex (green), nucleus (magenta), and ECM nodes (blue) are also shown. The simulation parameters correspond to *T* > *E* > *N* in Table 1.

Thus, hydrodynamics and incompressibility of the fluid *aid* migration in mechanism 2. The opposite scenario is observed for the push-pull mechanism of motility, where the pressure is low in the cell rear and high at the front of the cell near the protrusion (see Fig. 5(a)). The fluid is compressed at the front of the cell due to the cortical deformation at the leading protrusion as well as from the deformation of the ECM node. In spite of this adverse pressure gradient, the horizontal component of the cell’s velocity (as well as that of the fluid) is positive (Fig. 5(a), bottom) so that the cell is migrating from left to right. Stokes’ equations (see Eq. (S1.1) in Section S1, supporting text) are a force balance so that the restoring force of the ECM overcomes the pressure gradient force, enabling the cell to migrate. A simple analogy is to consider a rubber band that is stretched in a direction parallel to a pressure-driven background flow. Once the rubber band is sufficiently stretched and released, it will move in a direction opposite the fluid flow (from low to high pressure), dragging fluid with it. Fig. 5 shows that the ECM acts as this kind of rubber band in the push-pull mechanism of motility.

## Discussion

In this study, we have considered two mechanisms of cell migration: a mesenchymal-like push-pull and an amoeboid-like rear-squeezing mechanism. Three primary mechanical parameters of the model are the elasticities of the ECM and nucleus and the contractile tension of the cortex. The elasticities of the ECM and nucleus determine characteristic forces of the stretching of the ECM gap to the size of the undeformed nucleus, and of squeezing the nucleus to the width of the undeformed ECM gap, respectively. We find that the relation between three forces – the cortex’s contractile force and the characteristic forces of ECM and nucleus deformations – determine effectiveness of the cell migration for densely packed ECMs where the gap size is smaller than the nuclear diameter. The cell migrates most effectively when the contractile force exceeds both characteristic deformation forces, but the cell can also migrate persistently even if its tension is too weak to deform a near-rigid ECM, as long as the cortex’s contraction is sufficient to deform the nucleus. The cell can also migrate even if it fails to deform a stiff nucleus, but then the cell has to be able to sufficiently deform the ECM. Thus, the intuitive conclusion is that the cell has to contract enough to deform either ECM or nucleus, but not necessarily both. A nontrivial conclusion of our study is that a stiff nucleus limits the cell migration more than a stiff ECM does. Indeed, in the latter case the deformed nucleus is being squeezed from one ‘pocket’ of the ECM into the next one, while in the former case, the ECM is deformed around the stiff nucleus. In some cases the ECM relaxes back, instead of shifting the nucleus from one pocket to the next.

Another nontrivial prediction is that the rear-squeezing motility performs better than the push-pull mechanism in that the distance traveled per cycle is larger and experiences less variation across different parameter regimes. One reason for this prediction is the geometry of the model: in the push-pull mode, it is more likely, but not necessarily given, that the protrusion is made to the nearest node in front of the cell. The cell is then pulled to that node, and in a dense ECM network, the distance between the cell and the node is smaller than the cell diameter. In the rear-squeezing mode though, the cell is squeezed forward in between two nodes that are closer originally to the cell front, and so the cell often advances almost one body length. This prediction has to be used with caution: in reality, the protrusion could be made to a far-away node, in which case the push-pull mode would become more effective. Conversely, the rear-squeezing mechanism could be less effective if two random attachments to ECM nodes were used. Suppose in particular that the cell attempted to use two non-adjacent nodes to pass through a “gap” containing another node. In this case, the elasticity of the center node would block the cell from moving, since at some point the elastic force on the center node will push against the motion of the cell. In this study, we excluded these cases by mandating that protrusions be 30-45° apart on the cortex contour. Another relevant note is that we only investigated the mechanical constraints for the migration, not kinetic ones, as we did not examine how the speed of migration varies with changing parameters. In this setting a sparse ECM would fare poorly since the cell migration cycles would be prolonged by making protrusions that fail to bind to any nodes.

An important feature of our model is the underlying fluid and hydrodynamic forces. The hydrodynamic forces and movements effectively determine the interaction between the nuclear and cortical boundaries, as well as the interaction of each of these with the ECM. For example, in the rear-squeezing mode, a higher tension is required because the contraction at the rear must overcome the elasticity of the nucleus, which resists the contraction of the cortex through the hydrodynamic forces. Furthermore, because of the hydrodynamic forces, when the cell is stuck in the ECM, it truly is stuck, as any motion of the cell must carry the ECM with it due to the underlying fluid. The cell cannot slip away from the ECM unless there is sufficient space between it and the nodes. In this sense, we effectively model the parameter regimes where the cell is sufficiently loose to stay away from the ECM. Also, many previous studies have used artificial viscous dashpots to account for the movements and deformations generated by the elastic and contractile forces. In our case, the movements and deformations are mediated by actual viscous fluid. As a consequence, the presence of the fluid determines a timescale of the migration via a physical parameter, the viscosity, which can be measured experimentally. Lastly, our approach allows simulating the fluid pressure, the significance of which is highlighted by the prediction of the importance of nucleoplasmic pressure in pushing the nucleus through the ECM [35]. In our study, we determined that a favorable pressure gradient aids migration for cells using a rear-squeezing mechanism. We showed that intracellular pressure is high in the cell rear and low in the cell front so that the hydrodynamic contribution from fluid incompressibility aids migration. In contrast, we observe an adverse pressure gradient for the push-pull mechanism of migration.

The conclusions of our model are in qualitative agreement with a number of experimental studies. For example, dendritic cells of the immune system have soft and deformable nuclei, allowing them to move rapidly, a few microns per minute, in 3D [16]. By contrast, fibroblasts have stiffer nuclei and move through ECM slower than one micron per minute [4]. Cells reportedly move faster in softer ECM [12]. Cells were observed to move slower or even get stuck in conditions with stiffer nucleus or smaller ECM mesh size [49]. In fact, the problem of nuclear deformation is so dramatic that cells are known to use actomyosin cytoskeletal networks to actively deform the nucleus when migrating through narrow gaps [50]. Cells even use the elastic nucleus as a gauge to measure pore sizes and go to the least resistance path [51]. Last, but not least, one report supports the predicted advantage of the rear-squeezing mechanism: when neutrophils migrated through small holes, actin was seen to concentrate mostly at the cell rear, and myosin contraction was necessary to propel the cells through the holes [50].

Our simple model has, of course, a number of limitations. The main one is that we investigated a 2D caricature of the 3D geometry; the full 3D simulation would be prohibitively computationally expensive for multiple parameter sweeps. The structure of the cell in the model is simplified; for example, the model does not include mechanical connections between the lamin-based nucleoskeleton and the cytoskeleton, which facilitate force transmission between the nucleus and the extracellular matrix [52]. We have not explored in detail dependence of the motile behavior on mesh size and adhesion characteristics. The ECM, cortex and nucleus in our model are a linear elastic network, while the mechanical properties of actual ECM, cortex and nucleus are very complex [32, 53]. Needless to say, we have not investigated how cells change migration mode in response to the ECM properties [54], and we have not considered other modes of motility, such as blebbing [30] and chimneying [55]. More advanced problems, like mechanosensing [56], ECM remodeling and its feedback with cell mechanics [57], coordination of traction forces with ECM stiffness [11], proteolytic activity to melt down and rarefy the ECM in front of the cell [2, 5, 58], and nuclear damage in passages through small ECM gaps [59] are beyond the scope of the model.

The great modern surge in experimental research on 3D cell migration [4] is, fittingly, accompanied by modelling studies that started to address relevant mechanical questions [26, 54]. Very recently, modelling and experiment on 3D migration started to merge [60–62]. Our study, hopefully, will contribute to understanding of the general mechanical principles of 3D motility.

## Supporting information

Mechanism 1 through sparse ECM

Failure for mechanism 1

Mechanism 2 through sparse ECM

Failure for mechanism 2

Nuclear buckling and relaxation

## Supporting information

**S1 Text.** This file contains three sections. Section S1 describes the fluid dynamics and numerical methods used to solve the model equations. The re-meshing algorithm used to simulate mechanism 2 is described in Section S2. Section S3 contains a total of 11 figures showing the cell and ECM position at several time values for the parameter regimes listed in Table 1.

**S1 Video. Mechanism 1 through sparse ECM.** The cell migrates through a sparse ECM using mechanism 1 without deforming its nucleus.

**S2 Video. Failure for mechanism 1.** The cell becomes lodged in the ECM in simulations of the parameter regimes *E* > *N* > *T* and *N* > *E* > *T* using mechanism 1.

**S3 Video. Mechanism 2 through sparse ECM.** The cell migrates through a sparse ECM using mechanism 2 without deforming its nucleus.

**S4 Video. Failure for mechanism 2.** The cell becomes stuck in the ECM for parameter regimes *E* > *N* > *T* and *N* > *E* > *T* while migrating using mechanism 2.

**S5 Video. Nuclear buckling and relaxation.** A simulation of the parameter regime *T* > *N* > *E* (using mechanism 2) shows the cell nucleus wrinkles, or buckles, under high tension in the rear. After the cell detaches from the ECM nodes, the nucleus relaxes, which induces a flow that inhibits the cell’s forward progress through the ECM.

## Acknowledgements

We thank C. Copos for helpful discussions on the re-meshing algorithm. This work was supported by a grant from the Simons Foundation (#429808, Wanda Strychalski) and by US Army Research Office grant (#W911NF-17-1-0417, Alex Mogilner). Ondrej Maxian is supported by the NSF Graduate Research Fellowship DGE-1342536 and the Henry MacCracken fellowship.

## Author Contributions

**Conceptualization:** Ondrej Maxian, Wanda Strychalski, Alex Mogilner

**Data curation:** Ondrej Maxian, Wanda Strychalski

**Formal analysis:** Ondrej Maxian, Wanda Strychalski, Alex Mogilner

**Investigation:** Ondrej Maxian, Wanda Strychalski

**Methodology:** Ondrej Maxian, Wanda Strychalski, Alex Mogilner

**Project administration:** Wanda Strychalski, Alex Mogilner

