## Supporting text for "Computational estimates of mechanical constraints on cell migration through the extracellular matrix"

### S1. Fluid dynamics

For cellular motility problems, small length scales (microns) and slow movements (microns per minute) lead to extremely small Reynolds numbers [1], and the fluid permeating the cell and ECM obeys the Stokes' equations:

$$\mu \Delta \boldsymbol{u} - \nabla p = -\boldsymbol{f} \tag{S1.1}$$

$$\nabla \cdot \boldsymbol{u} = 0. \tag{S1.2}$$

Here $\mu$ is the fluid viscosity, $p$ is the pressure, $\boldsymbol{u}$ is the fluid velocity, and $\boldsymbol{f}$ is an external force (which comes from any immersed solid structures). For simplicity, we assume that the nucleoplasm, cytoplasm and interstitial fluids all have the same viscosity.

#### *S1.1. Solution method: regularized Stokeslets*

Because the Stokes' equations are linear, boundary integral methods can be used to solve the system in Eqs. (S1.1) and (S1.2). In particular, the solution to the Stokes' equations in free space can be determined by convolving the external force function $\boldsymbol{f}$ against the free space Green's function for Stokes' flow over the interface or immersed structure where the force is applied [2]. However, the singularity of the Green's function at the point where the force is applied must be treated with accurate quadrature rules, or the singular integral must be regularized. We choose to regularize the problem with the method of regularized Stokeslets [3]. In particular, we suppose that the external force of strength $\boldsymbol{f}_0$ is distributed over a small ball centered on a point $\boldsymbol{x}_0$ (rather than confined exclusively to the point $\boldsymbol{x}_0$), so that

$$\boldsymbol{f}(\boldsymbol{x}) = \boldsymbol{f}_0 \phi_\epsilon(\boldsymbol{x} - \boldsymbol{x}_0). \tag{S1.3}$$

The solution to the Stokes' equations with external forcing given by Eq. (S1.3) can be derived from the form of the "blob" or "cutoff" function $\phi_\epsilon$. For example, if

$$\phi_\epsilon(\boldsymbol{x}) = \frac{3\epsilon^3}{2\pi(\|\boldsymbol{x}\|^2 + \epsilon^2)^{5/2}}, \tag{S1.4}$$

then

$$p^\epsilon(\boldsymbol{x}, \boldsymbol{x}_0) = \frac{1}{2\pi}(\boldsymbol{f}_0 \cdot (\boldsymbol{x} - \boldsymbol{x}_0)) \left( \frac{r_0^2 + 2\epsilon^2 + \epsilon\sqrt{r_0^2 + \epsilon^2}}{(\sqrt{r_0^2 + \epsilon^2} + \epsilon)(r_0^2 + \epsilon^2)^{3/2}} \right), \tag{S1.5}$$

and

$$\boldsymbol{u}^\epsilon(\boldsymbol{x}, \boldsymbol{x}_0) = -\frac{\boldsymbol{f}_0}{4\pi\mu} \left( \ln\left( \sqrt{r_0^2 + \epsilon^2} + \epsilon \right) - \frac{\epsilon(\sqrt{r_0^2 + \epsilon^2} + 2\epsilon)}{(\sqrt{r_0^2 + \epsilon^2} + \epsilon)\sqrt{r_0^2 + \epsilon^2}} \right) + \frac{1}{4\pi\mu}(\boldsymbol{f}_0 \cdot (\boldsymbol{x} - \boldsymbol{x}_0))(\boldsymbol{x} - \boldsymbol{x}_0) \frac{\sqrt{r_0^2 + \epsilon^2} + 2\epsilon}{(\sqrt{r_0^2 + \epsilon^2} + \epsilon)^2 \sqrt{r_0^2 + \epsilon^2}} \tag{S1.6}$$

are the pressure and velocity that result from the force in Eq. (S1.3), where $r_0 = \|\boldsymbol{x} - \boldsymbol{x}_0\|$. The derivation of these expressions can be found in [3]. An equivalent formulation is to write the result of Eq. (S1.6) as a matrix-vector product (since the system is linear), so that

$$\boldsymbol{u}(\boldsymbol{x}) = \mathbf{M}(\boldsymbol{x}, \boldsymbol{x}_0)\boldsymbol{f}(\boldsymbol{x}_0), \tag{S1.7}$$

where the matrix $\mathbf{M}$ is determined from the locations of the points $\boldsymbol{x}$ and $\boldsymbol{x}_0$.

The key point here is that given the external forcing term(s) $\boldsymbol{f}_0$, the velocity at any point $\boldsymbol{x}$ in the domain can be determined by Eq. (S1.7). We note that the units in Eq. (S1.6) imply that $\boldsymbol{f}_0$ has units pN/$\mu$m.

*S1.2. Stokes' paradox and force imbalances in 2D*

A complication arises when modeling objects that have net nonzero forcing in 2D. Here we wish to consider an extracellular matrix (ECM) composed of fibers that are tethered together by springs. In that case, it is virtually impossible for the net force on the ECM to be zero during a dynamic simulation, where by definition the fibers are moving and the resulting forces are changing. A net force on the domain inevitably results from our model of the ECM. The fundamental solutions of Stokes' flow in free space (regularized in Eq. (S1.6)) contain a logarithmic term so that net force imbalances result in velocities as $||\boldsymbol{x}|| \to \infty$. Thus the "usual" free space boundary condition of zero velocity at $\infty$ is incompatible with the net forcing, and a new boundary condition must be imposed [4]. One option is to specify a rigid ECM, which leads to a no-flow boundary condition on the surface of the ECM fibers. Here we study the effect of ECM elasticity and want to account for fiber flexibility. We therefore follow our previous work [5] and impose the boundary condition of *zero mean flow on a large circle of radius R that surrounds the domain.* As $R \to \infty$, this boundary condition is equivalent to the usual free space boundary condition. We use the value of $R = 1000$ (cell diameters) as in [5].

The boundary condition on the large circle of radius $R$ allows for net nonzero forcing within the domain and can be enforced by adding a constant velocity throughout the domain of flow. In [5], we derived this velocity for a single point force as

$$\boldsymbol{u}^R(\boldsymbol{f}_0) = -\frac{\boldsymbol{f}_0}{4\pi\mu}\left(\frac{1}{2} - \ln R\right). \tag{S1.8}$$

If there are $N$ points where force is applied, each with force $\boldsymbol{f}_k$, the constant velocity added throughout the domain is

$$\boldsymbol{u}^{\boldsymbol{R}} = \sum_{k=1}^{N} \boldsymbol{u}^R(\boldsymbol{f}_k) = -\sum_{k=1}^{N} \frac{\boldsymbol{f}_k}{4\pi\mu}\left(\frac{1}{2} - \ln R\right). \tag{S1.9}$$

This ensures that the additional boundary condition is satisfied and yields a well-posed problem.

The full numerical scheme is therefore as follows: given a collection of $N$ points (discretized fibers) with applied forces $\boldsymbol{f}_k$, Eq. (S1.7) is used to compute an intermediate velocity at each of the $N$ points. Then the boundary condition is enforced through the addition of a constant velocity in Eq. (S1.9), so that the total fluid velocity at each point is the sum of the two. Explicit first-order time-stepping is then used to update the positions of the points.

One of the numerical parameters in the model is $\epsilon$, the width of the blob function in hydrodynamic Eq. (S1.4). As discussed in [3] and [6], $\epsilon$ should be on the order of the spacing of the grid in the system. We set $\epsilon = 0.075$ cell diameters.

### S2. Re-meshing algorithm

Here we test our re-meshing algorithm outside of the cell motility simulations. We consider a closed elastic fiber of radius 1 discretized with $N = 50$ points, sitting by itself in a domain of fluid of viscosity $\mu = 1$ Pa·s inside and outside. The back half of the cell (points $\lceil N/4 \rceil$ to $\lfloor 3N/4 \rfloor$) is stiff with $k = 100$ pN/$\mu$m and resting spring length $r = 0.5$, while the front of the cell (blue circles) is loose, with $k = 1$ pN/$\mu$m and $r = 1$. These parameters are similar as those used in the mechanism 2 simulations, with the exception that we choose here to keep $r = 0.5$ so that the dynamics of the back of the cell are similar to those shown in mechanism 2 simulations.

We simulate the dynamics of the circle within the method of regularized Stokeslets with $\epsilon$ equal to the initial point spacing. We begin by simulating the cell to steady state ($t = 7$ with $\Delta t = 0.01$)

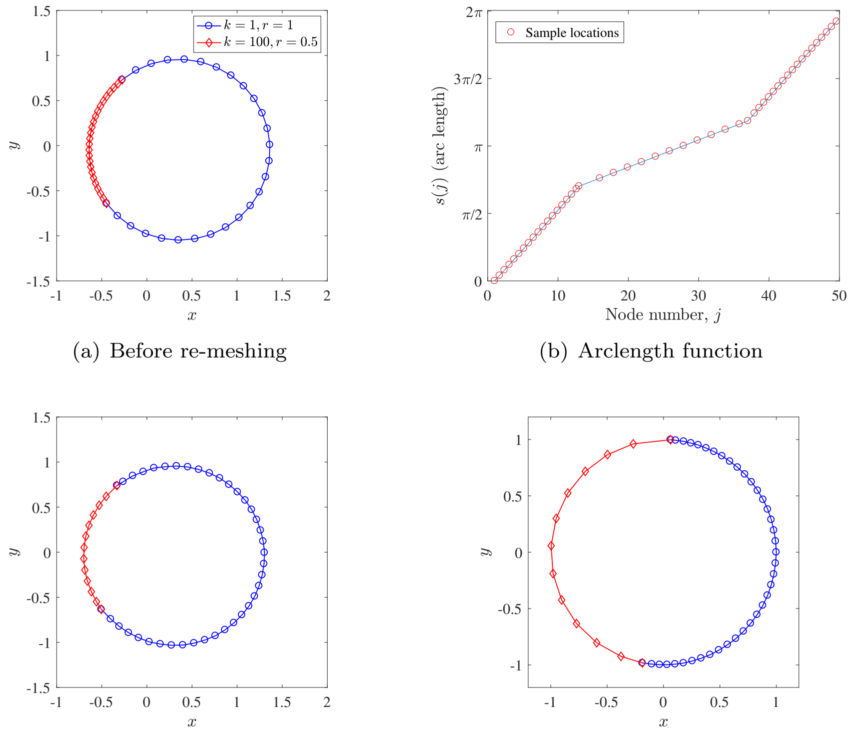


(a) Before re-meshing (b) Arclength function

(c) After re-meshing (d) New reference locations

Figure S1: Test on the re-meshing algorithm. (a) Cell before re-meshing. The back of the cell (red diamonds) is rigid with $k = 100$ pN/$\mu$m and $r = 0.5$, while the front of the cell (blue circles) has $k = 1$ pN/$\mu$m and $r = 1$. (b) The arc length function $s(j)$ is sampled at equal distances to give the new node locations, which are interpolated from the old locations. (c) After re-mesh. We note how the points are now evenly distributed. The point of interest in the error calculation is the first counter-clockwise red point in the second quadrant of the cell. (d) The new reference locations used to calculate the elastic force after re-meshing.

without re-meshing. Fig. S1(a) shows the deformed shape at this point. We note how the original back half of the cell (points $\lceil N/4 \rceil$ to $\lfloor 3N/4 \rfloor$ marked with red diamonds) has collapsed down, leaving the front-half potentially unresolved. Fig. S1(b) shows the corresponding arclength function $s(j)$ at this point in the algorithm as a blue line. We note how from indices 13 to 37, ($\lceil N/4 \rceil$ to $\lfloor 3N/4 \rfloor$) the slope of the arclength function changes drastically because the points are more tightly packed. In Fig. S1(b), the new *sample locations* are shown as red circles. We note how these locations are more sparse in the region of decreased slope. This indicates that the arclength function $s(j)$ is being sampled at equally spaced node indices, $S(j)$. Indeed, as shown in Fig. S1(c), the points $\boldsymbol{X}(S(j))$ are equally spaced on the cell after they are re-meshed (we always keep the 2 points bound to the ECM nodes). Finally, Fig. S1(d) shows the new reference configurations, $\boldsymbol{X}^0(S(j))$. We note how these become more sparse at the back of the cell.

The re-mesher always includes the two points bound to ECM nodes. Because of this, an ideal test for the re-mesher is to consider the displacement of one of these points with and without re-meshing. That is, we vary the number $N$ of points on the cortex and simulate to $t = 7$ (empirically determined to be steady state) with $\Delta t = 0.01$ as before. Next, we simulate to $t = 8$ without and with re-meshing (only once at $t = 7$) and determine how much the first red point in the second quadrant (moving counterclockwise) has moved relative to not re-meshing (i.e. the error is the norm of the position of the point at $t = 8$ with re-meshing minus the position without re-meshing). Table S1 shows the results of this test. It appears that our re-mesher makes an error on the order $\mathcal{O}(10^{-4})$ for $\approx 100$ points, which is what we use in our simulations. This error could be decreased

| $n$ | 25 | 50 | 100 | 200 |
|---|---|---|---|---|
| Error | $5.5 \times 10^{-3}$ | $1.6 \times 10^{-3}$ | $4.1 \times 10^{-4}$ | $1.2 \times 10^{-4}$ |

Table S1: Errors in the final position of the first red node in the second quadrant (in counterclockwise order).

by increasing the number of points.

We note that the procedure for re-meshing the nucleus is exactly the same as that described for the cortex, with the exception that there are no "bound points" that must be preserved by the re-mesher. Thus the sampling of the arclength function is truly uniform in this case.

### S3. Supporting Figures

The figures in this section show the position of the cortex (green), nucleus (magenta), and ECM nodes (blue) at several time values during a simulation. When simulations show significant deformation of the ECM, the initial position of the nodes is indicated by black ×'s. Time values increase from left to right and top to bottom for figures in this section. Key parameter values for each simulation are indicated in the figure captions using bold text. See Table 1 in the main text for exact parameter values.

#### *S3.1. Push-pull mechanism of motility*

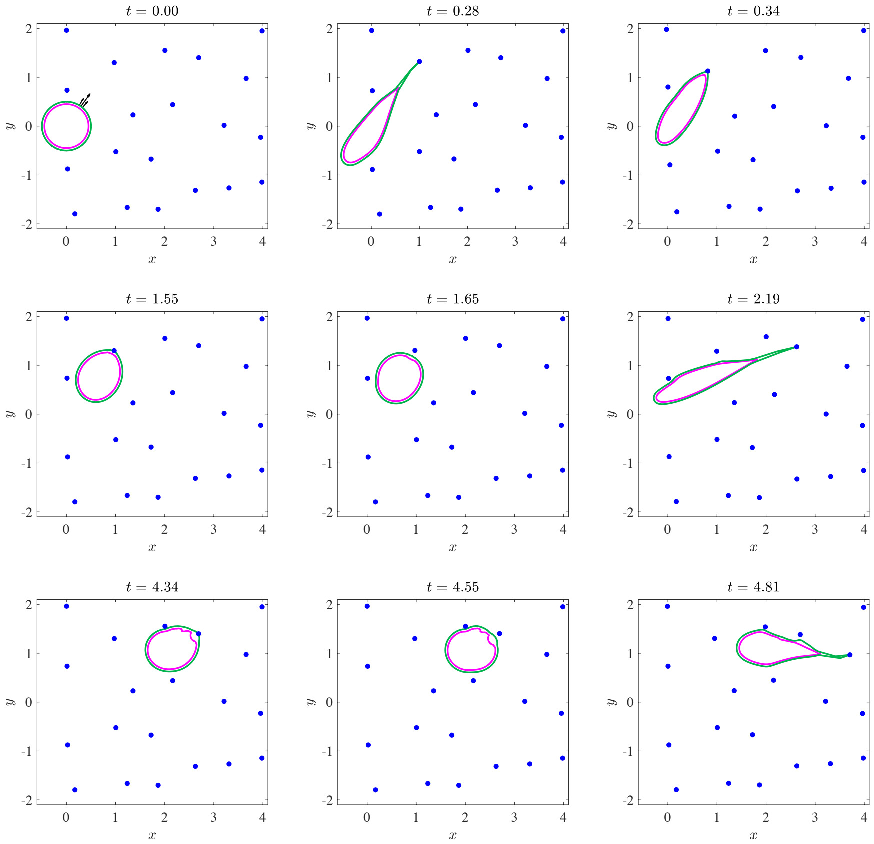


Figure S2: **Sparse ECM**: The cell migrates easily through the sparse network (same as the network shown in Fig. 1(a) of the main text).

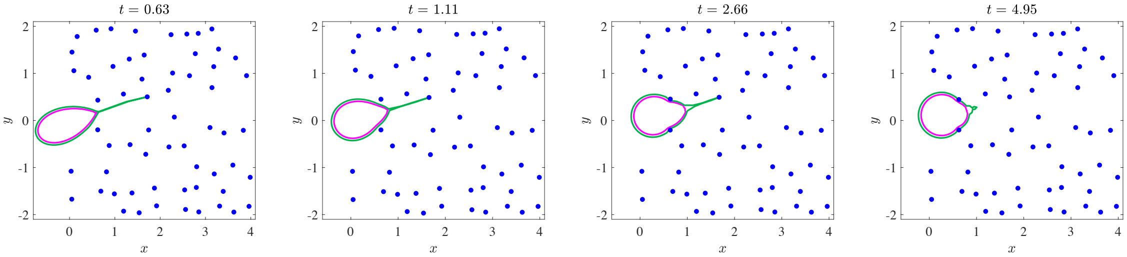


Figure S3: $\boldsymbol{E} > \boldsymbol{T}$ **and** $\boldsymbol{N} > \boldsymbol{T}$. The cell becomes stuck in the ECM during the simulation.

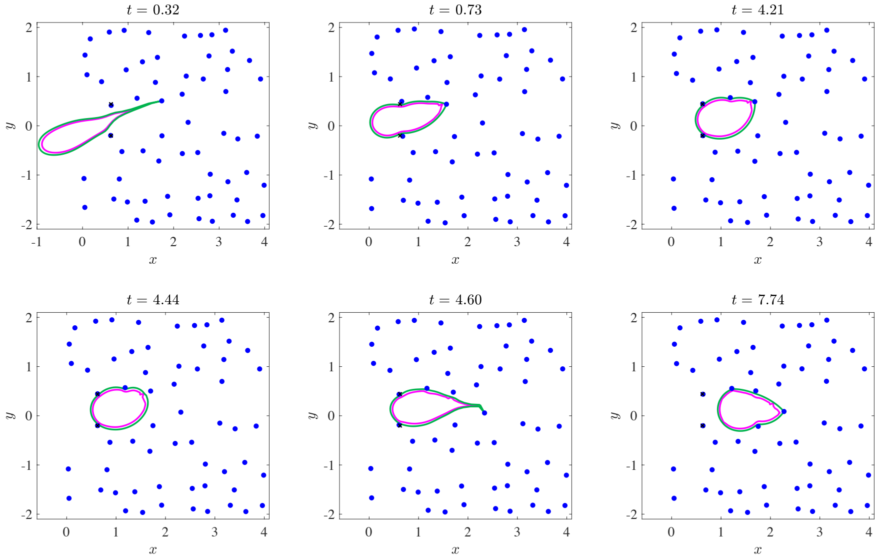


Figure S4: $\boldsymbol{T} > \boldsymbol{E} > \boldsymbol{N}$: The cell migrates by squeezing the nucleus, which is relatively more deformable than the cortex and ECM.

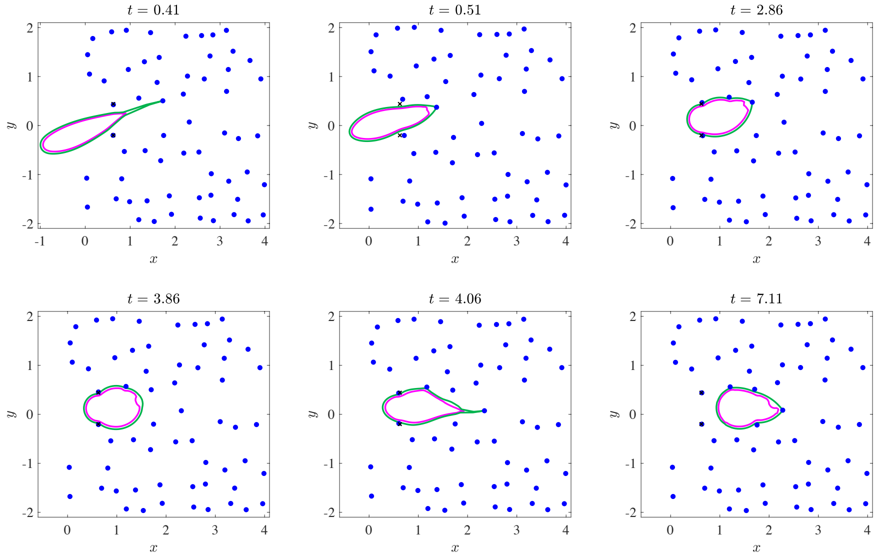


Figure S5: $\boldsymbol{T} > \boldsymbol{N} > \boldsymbol{E}$. The cell moves using a combination of nuclear and ECM deformation. Black $\times$'s indicate the original position of the key ECM nodes.

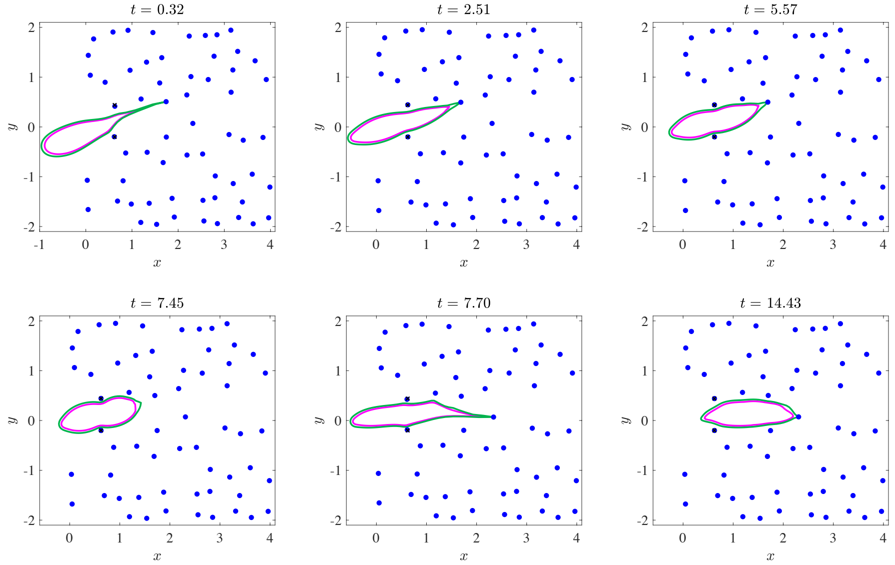


Figure S6: $\boldsymbol{E} > \boldsymbol{T} > \boldsymbol{N}$: The cell moves by squeezing the nucleus because of relatively reduced tension in the cortex. Note the elongated cell shape for this parameter regime.

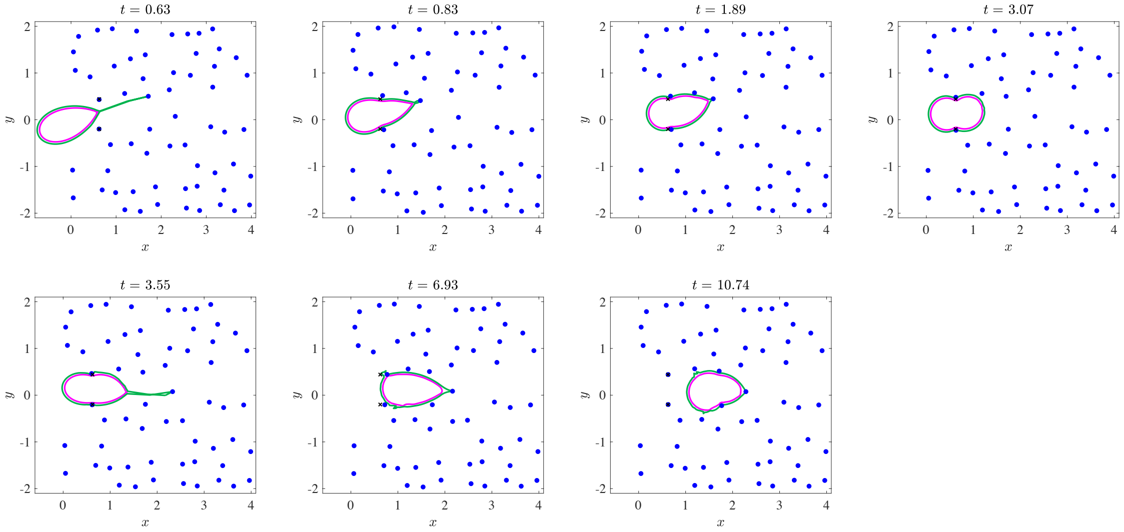


Figure S7: $\boldsymbol{N} > \boldsymbol{T} > \boldsymbol{E}$: The cell moves by a combination of nuclear and ECM deformation. However, in this paramater regine, the cell must deform the ECM to migrate because the nucleus is relatively stiffer (compare to the data in Fig. S6). Black $\times$'s are used to show the initial position of the key ECM nodes, which achieve their maximum displacement at $t = 6.93$.

*S3.2. Rear-squeezing mechanism of motility*

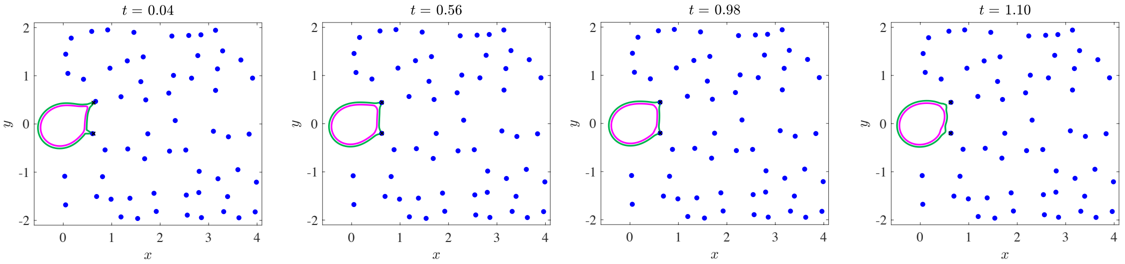


Figure S8: $\boldsymbol{E} > \boldsymbol{T}$ **and** $\boldsymbol{N} > \boldsymbol{T}$: When the contraction of the cell cortex is too weak to deform either the nucleus or ECM, the cell is not able to squeeze through a gap in the ECM.

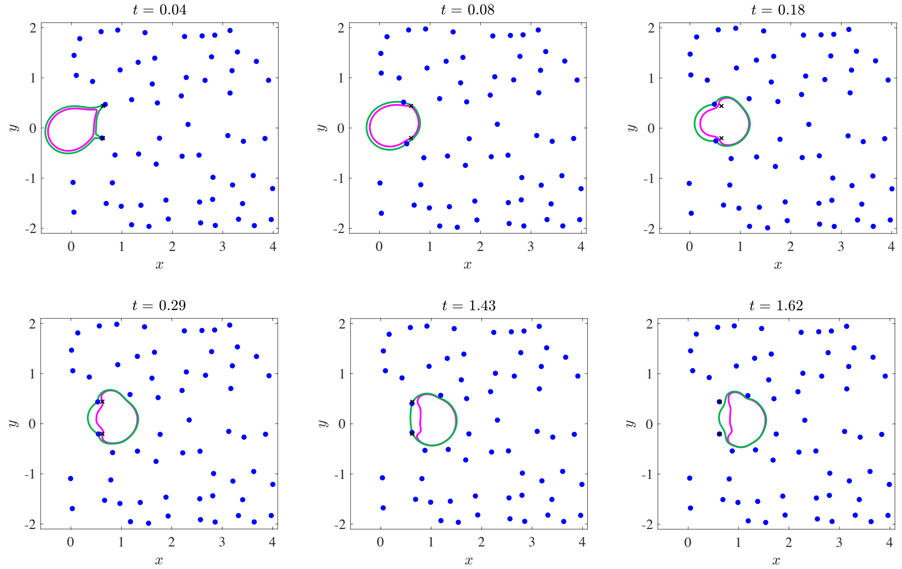


Figure S9: $\boldsymbol{T} > \boldsymbol{E} > \boldsymbol{N}$: The cell can easily pass through the gap between the fibers because the nucleus is soft compared to the cortex and ECM.

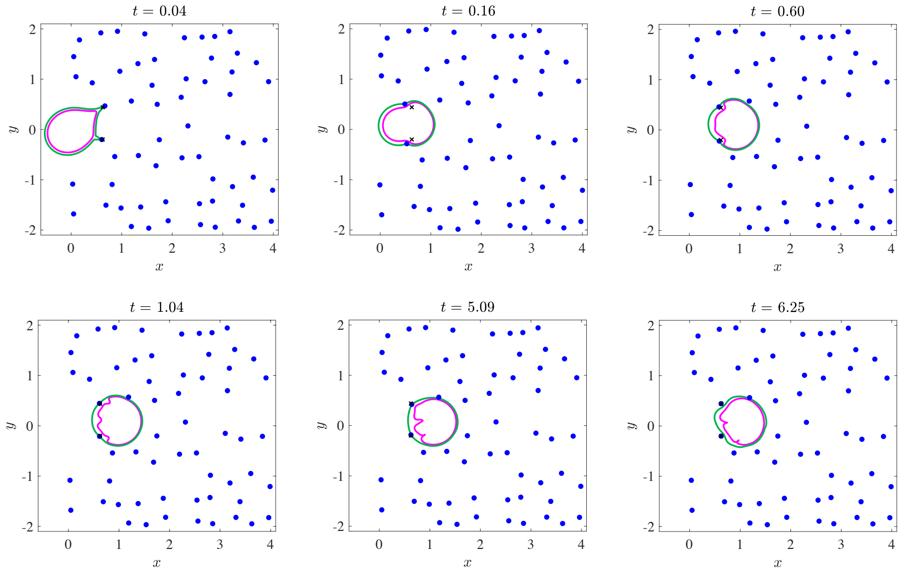


Figure S10: $\boldsymbol{T} > \boldsymbol{N} > \boldsymbol{E}$: The nucleus begins to buckle (bottom row) because of high cortical tension and increased cortical stiffness. After releasing the nodes, the nucleus partially regains its rounded shape (compare the position of the nucleus at $t = 6.25$ to $t = 5.09$).

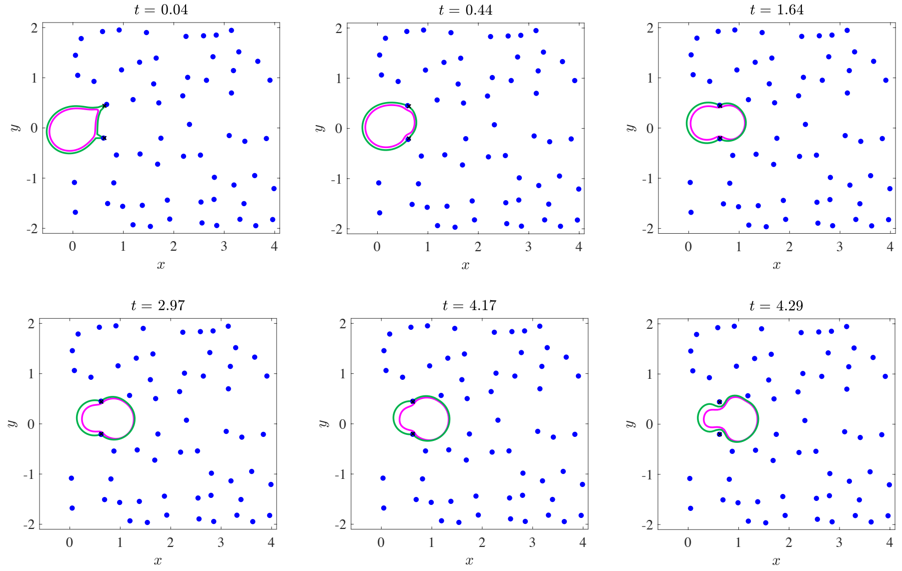


Figure S11: $\boldsymbol{E} > \boldsymbol{T} > \boldsymbol{N}$: Although the cell makes some progress, there is not enough force to drive it completely through the gap. After one motility cycle, the cell retracts slightly after releasing the nodes.

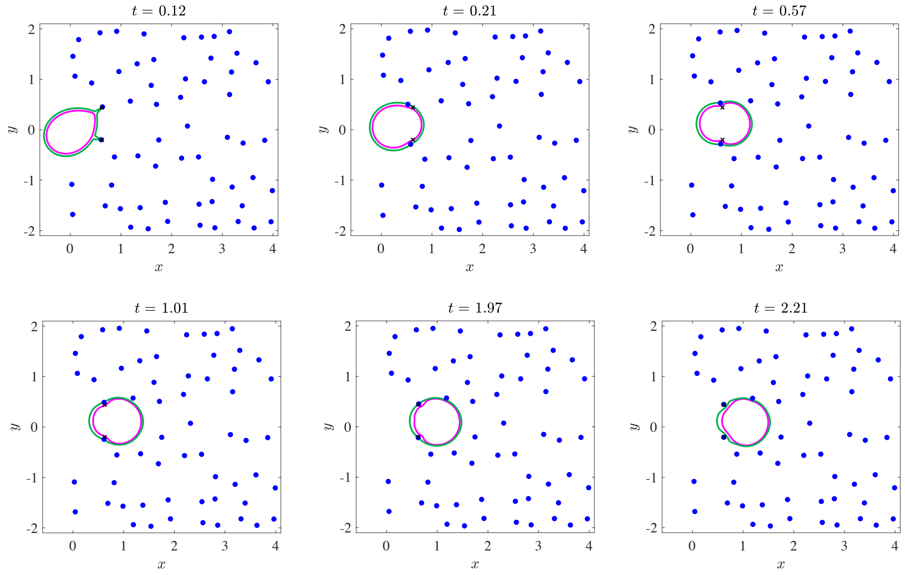


Figure S12: $\boldsymbol{N} > \boldsymbol{T} > \boldsymbol{E}$: In spite of the nucleus' relatively large stiffness, the cell is still able to migrate through the ECM. However, the cell is unable to entirely pass through the gap after one motility cycle.
